# Humanized tauopathy chimeras uncover microglial and lncRNA strategies for neuroprotection

**DOI:** 10.64898/2026.07.02.736124

**Authors:** Wenhui Qu, Li Fan, Minwoo Wendy Jang, Pearly Ye, Ethan Cordes, Abulimiti Aikedan, Wen Hu, Ravi Kumar Nagiri, Man Ying Wong, Wenjie Luo, Mathew Blurton-Jones, Hagen U Tilgner, Anna Orr, Shiaoching Gong, Li Gan

## Abstract

Human genetics implicates innate immunity as a key modifier of tau toxicity, yet human-specific neuroimmune mechanisms remain difficult to test in vivo. Here, we developed HuMiNAX, the first humanized iPSC-based neuroimmune xenograft model of tau-associated neurodegeneration, enabling human microglia to interact with human neurons and astrocytes in the adult mouse brain. In HuMiNAX, tau seeding induced aggregation only in mutation-carrying human neural grafts, causing neuron loss and inflammatory activation of human microglia. Progranulin-overexpressing human microglia dampened tau-associated inflammation, preserved neurons, and restored neuronal gene-expression and RNA-splicing programs, supporting microglial control of neuronal resilience. CRISPRi knockdown of the human-specific lncRNA HNRNPK-AS1 also protected neurons in HuMiNAX. These findings establish HuMiNAX as a human neuroimmune model of tauopathy and identify microglial and RNA-mediated strategies of neuronal resilience.

---

A fundamental question in neurodegenerative disease research is how the accumulation of misfolded proteins is translated into progressive neurodegeneration (*1*). Although protein aggregation is a defining pathological feature across disorders, its relationship to neuronal loss is neither uniform nor fully deterministic, implying that additional cellular programs govern neuronal vulnerability (*2, 3*). Large-scale human genetics has provided critical insight into these programs, revealing that Alzheimer’s disease (AD) risk variants are strikingly enriched in genes expressed by microglia, implicating innate immune pathways as central modulators of disease progression rather than passive bystanders (*4*).

Tauopathies, including AD and related dementias, provide a compelling context in which to address this problem, as the spread of pathological tau aggregation is tightly linked to disease progression and cognitive decline (*5*). Consistent with genetic evidence, growing functional studies demonstrate that microglia actively regulate tau propagation and toxicity (*6*). However, increasing evidence highlights pronounced species differences in disease-associated transcriptional programs, particularly within microglia, where human and mouse cells can mount divergent responses to tau pathology (*7–9*). In addition, human-specific regulatory architectures, including extensive networks of long noncoding RNAs (lncRNAs), are poorly conserved in rodents, constraining the ability of mouse models to capture human-specific pathways that may critically shape neuronal vulnerability (*10, 11*). Together, these observations underscore the need for human-relevant systems that enable mechanistic interrogation of microglial gene networks, tau propagation, and neurodegeneration in a genetically and transcriptionally faithful context (*12, 13*).

Human-induced pluripotent stem cell (iPSC)-derived brain cells provide a powerful framework for modeling neurodegenerative disease in a human context. Using 2D cultures, we previously established a robust tau seeding model in engineered human iPSC lines that express adult four repeat (4R) tau carrying the P301S mutation linked to frontotemporal dementia (FTD) (*14*). 3D organoid systems extend this approach by introducing cellular diversity, yet they remain limited by incomplete maturation and the absence of vascular integration (*15*). Xenotransplantation into the mouse brain overcomes these constraints by placing human iPSC-derived brain cells in a living tissue environment that supports their long-term survival, differentiation, functional maturation, and disease-relevant interactions in vivo (*16, 17*). However, prior transplantation studies have typically examined human neurons or human microglia in isolation (*18–22*). As a result, critical neuroimmune interactions that shape tau-induced neurodegeneration remain poorly defined.

Here, we establish HuMiNAX, a humanized neuroimmune xenograft platform that integrates human neurons, astrocytes, and microglia while replacing endogenous murine microglia. By combining human iPSC-derived brain cells that support robust tau propagation (*14*) with human microglia, HuMiNAX enables mechanistic analysis of tau-induced neurodegeneration in a human cellular context in vivo. Using single-nucleus and spatial transcriptomics, we show that tau propagation selectively compromises specific human neuronal populations and elicits a strong innate immune response in human microglia marked by interferon signaling and suppression of progranulin associated pathways.

Leveraging the perturbability of HuMiNAX, we set out to determine how microglial state influences neuronal vulnerability to tau pathology by selectively elevating progranulin signaling in human microglia. We further sought to identify neuron intrinsic mechanisms that act downstream of this microglial state and mediate its effects on neuronal survival and identified human-specific lncRNA HNRNPK-AS1 as a candidate RNA regulatory node linking progranulin-dependent microglial signaling to neuronal responses to tau aggregation. Together, this framework positions HuMiNAX as an in vivo platform to dissect how immune-driven states engage RNA regulatory programs to shape neuronal fate in neurological disorders.

## HuMiNAX supports integrated development of human neurons, astrocytes, and microglia in vivo

Neuroimmune interactions play critical roles in neurodegenerative disease, but perturbable in vivo systems composed of multiple human brain cell types remain limited. To address this gap, we established **HuMiNAX**, a human iPSC-derived **mi**croglia, **n**euron, and **a**strocyte **x**enograft platform designed to model human neuroimmune interactions in vivo. This platform builds on our previous neural xenograft work (*18*) by incorporating a published human microglia replacement strategy (*20*) and optimizing the transplantation paradigm to achieve consistent co-engraftment of human neural and microglial lineages in the adult mouse brain. To model adult 4R tauopathies, we first established 4R-WT NPC grafts, since conventional iPSC-derived neurons predominantly express fetal 3R tau and show limited expression of the adult-associated 4R tau isoforms central to tauopathies (*14*). Before transplantation, these cryopreservable NPCs expressed neural progenitor markers Nestin and SOX1 and the proliferation marker KI67 (**fig. S1A**).

To generate HuMiNAX, we co-transplanted equal numbers (105,000 cells) of 4R-WT NPCs and human HPCs carrying PLX5622-resistant CSF1R^G795A^ into the posterior hippocampus of 2-4 month-old young immunotolerant mice (hCSF1KI/KI; Rag2-/-; Il2rg-/-) after PLX5622-mediated depletion of endogenous mouse microglia (*20*). These mice express human CSF1 to support human microglia and lack mature T, B, and NK cells to permit long-term engraftment. PLX5622 treatment was maintained for an additional 1.5 months after transplantation to suppress mouse microglia while enabling CSF1R^G795A^ human microglia to differentiate and expand throughout the brain (*20*) (**Fig. 1A**).

**Fig. 1.**
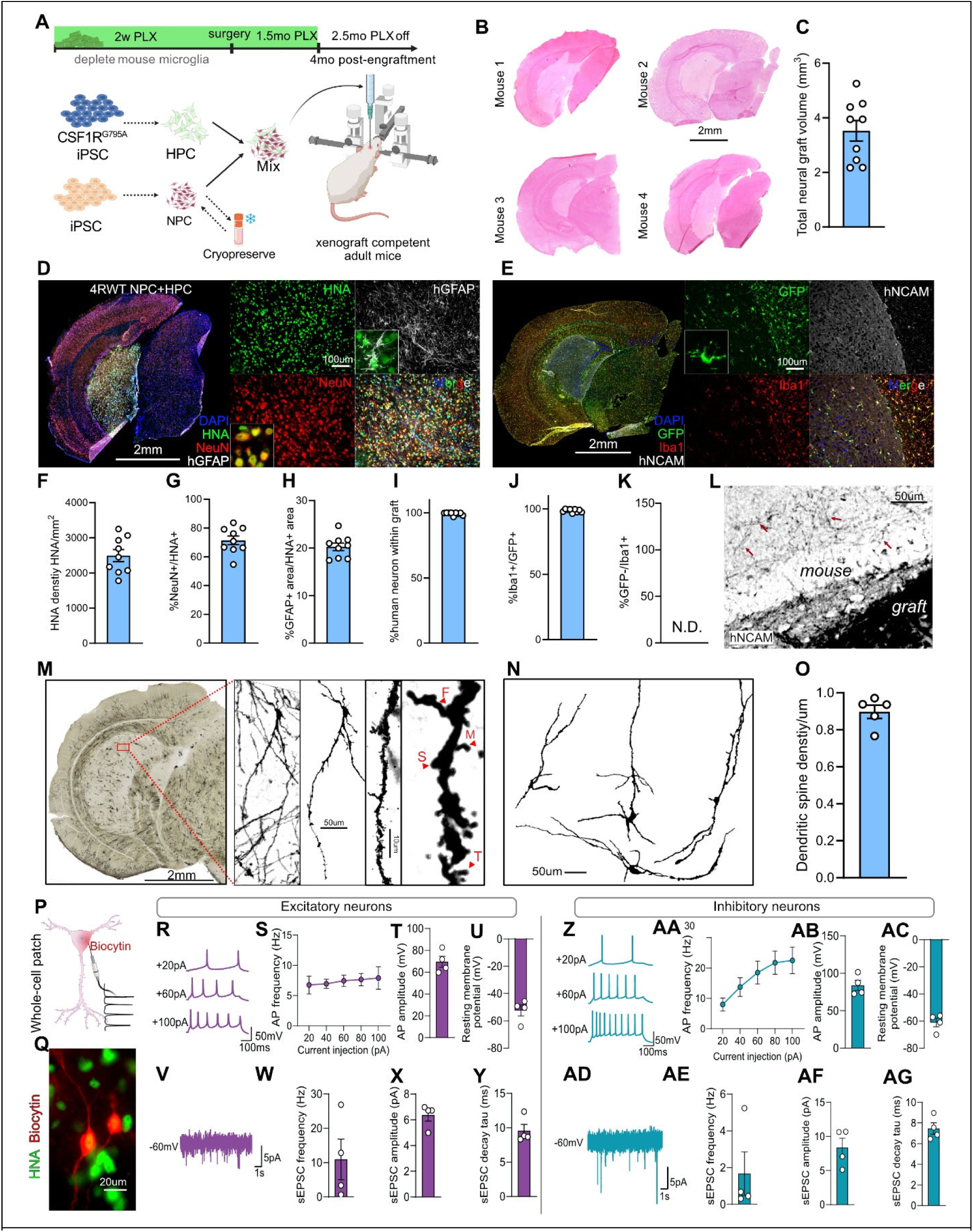
Establishment and functional characterization of the HuMiNAX human neuroimmune chimera platform. (**A**) Experimental schematic of HuMiNAX generation, including differentiation of CSF1R^G795A^ human iPSC derived hematopoietic progenitor cells (HPCs) and human iPSC derived neural progenitor cells (NPCs), PLX5622 treatment, stereotactic transplantation, and post engraftment analysis. (**B-C**) Representative coronal brain sections from transplanted mice (B) and quantification of total neural graft volume (C). N=9 mice. (**D**) Representative immunofluorescence images of human neural grafts stained for DAPI, HNA, NeuN, and hGFAP. (**E**) Representative immunofluorescence images of human microglia within hNCAM+ human neural grafts stained for DAPI, GFP, IBA1, and hNCAM. (**F-K**) Quantification of graft composition, including HNA+ cell density (F), percentage of HNA+ cells expressing NeuN (G), hGFAP+ area normalized to HNA+ graft area (H), percentage of human neurons within the graft (I), percentage of GFP+ cells expressing IBA1 (J), and percentage of GFP negative IBA1+ mouse microglia within the graft (K). N.D., not detected. N=9 mice. (**L**) Representative hNCAM staining at the graft border. (**M-O**) Golgi Cox staining of grafted human neurons (M), representative neuronal reconstruction (N), and quantification of dendritic spine density (O). F, filopodia; S, stubby; M, mushroom; T, thin. N=5 mice. (**P-Q**) Whole cell patch clamp recording schematic (P) and post hoc biocytin and HNA labeling of a recorded human neuron (Q). N=4 mice. (**R-U**) Representative evoked action potential traces from excitatory neurons (R) and quantification of action potential frequency (S), action potential amplitude (T), and resting membrane potential (U). N=4 mice. (**V-Y**) Representative spontaneous excitatory postsynaptic current recording from a graft derived excitatory neuron (V) and quantification of sEPSC frequency (W), amplitude (X), and decay tau (Y). N=4 mice. (**Z-AC**) Representative evoked action potential traces from inhibitory neurons (Z) and quantification of action potential frequency (AA), action potential amplitude (AB), and resting membrane potential (AC). N=4 mice. (**AD-AG**) Representative spontaneous excitatory postsynaptic current recording from graft derived inhibitory neurons (AD) and quantification of sEPSC frequency (AE), amplitude (AF), and decay tau (AG). Data are shown as mean ± SEM. For histological and electrophysiological quantifications, each data point represents the average value from one mouse.

GFP expression in transplanted human HPCs enabled identification of HPC-derived human microglia and their distinction from host microglia (*20*). GFP+ human microglia expanded within hNCAM+ human neural grafts between two and four months after engraftment, while CD31+ vascular networks became more continuous (**fig. S1B,C**). Based on the maturation timeline and prior evidence of FTD mutation-associated neuronal loss by four months post-engraftment (*23*), we selected four months for subsequent analyses, when HuMiNAX reproducibly generated sizable human neural grafts across animals (**Fig. 1B,C**). 4R-WT NPCs differentiated into hNA+ NeuN+ neurons and hGFAP+ astrocytes, while co-transplanted HPCs produced widespread GFP+ IBA1+ human microglia that populated hNCAM+ grafts and extended into host tissue (**Fig. 1D-E**). Quantification confirmed robust engraftment, neuronal and astrocytic differentiation, and complete replacement of mouse microglia within the graft, with no GFP- IBA1+ mouse microglia detected in the human graft area (**Fig. 1F-K**). hNCAM+ human neuronal processes also extended into surrounding mouse parenchyma, consistent with local outgrowth of graft-derived human neurites (*18*).

We next assessed neuronal maturation, which was not well defined in our prior NPC transplantation models (*18*). Golgi-Cox staining revealed graft-derived human neurons with elaborate dendritic arbors, abundant spines, and diverse spine morphologies, including mushroom, stubby, thin, and filopodia-like spines (**Fig. 1M**). Three-dimensional reconstructions and quantification confirmed complex neuronal morphology and robust spine density (**Fig. 1N,O; fig. S2A-D**). Whole-cell patch-clamp recordings further demonstrated functional maturation. Biocytin-filled recorded cells were confirmed as human by HNA immunostaining (**Fig. 1P,Q**). Excitatory neurons fired action potentials and showed spontaneous excitatory postsynaptic currents, indicating intrinsic excitability and excitatory synaptic input (**Fig. 1R-Y**). Inhibitory neurons likewise fired action potentials and displayed spontaneous excitatory postsynaptic currents, supporting functional maturation of inhibitory neuronal populations (**Fig. 1Z-AG**).

To determine whether HuMiNAX supports neuroimmune maturation across distinct human NPC lines and tau isoform contexts, we co-transplanted 3R-*MAPT* (3R-WT) NPCs with HPCs. Before transplantation, 3R NPCs also expressed Nestin, SOX1, and KI67 (**fig. S1D**). Similar to 4R-WT grafts, 3R NPC + HPC grafts produced HNA+ human neurons, hGFAP+ astrocytes, and GFP+ IBA1+ human microglia within hNCAM+ human neural tissue (**fig. S1A-F**). Graft-associated human microglia expressed P2RY12 and TMEM119, supporting acquisition of a homeostatic identity in vivo (**fig. S1G,I**). 3R-MAPT graft-derived neurons showed intrinsic excitability, spontaneous synaptic activity, dendritic arborization, and spine morphology comparable to 4R-WT neurons (**fig. S1J-Y; fig. S2E-I**). Thus, HuMiNAX reproducibly supports coordinated maturation of human neurons, astrocytes, and microglia across 3R and 4R-WT grafts.

## Single-nucleus profiling reveals diverse human neuronal subtypes and glial maturation in HuMiNAX

The large, spatially defined HuMiNAX grafts enabled microdissection and recovery of high-quality human nuclei for snRNA-seq. After quality control and dataset integration (*24*), we obtained 26,102 human nuclei from four 4R-WT NPC + HPC grafts (**Fig. 2A; fig. S3A-H**). Cell-type annotation was refined using canonical markers, cell-cycle analysis, and subclustering (**fig. S3I-K**). RNA velocity analysis revealed progressive differentiation of NPC-derived lineages into neuronal and astrocytic populations, alongside maturation of the HPC-derived microglial lineage (**Fig. 2B**).

**Fig. 2.**
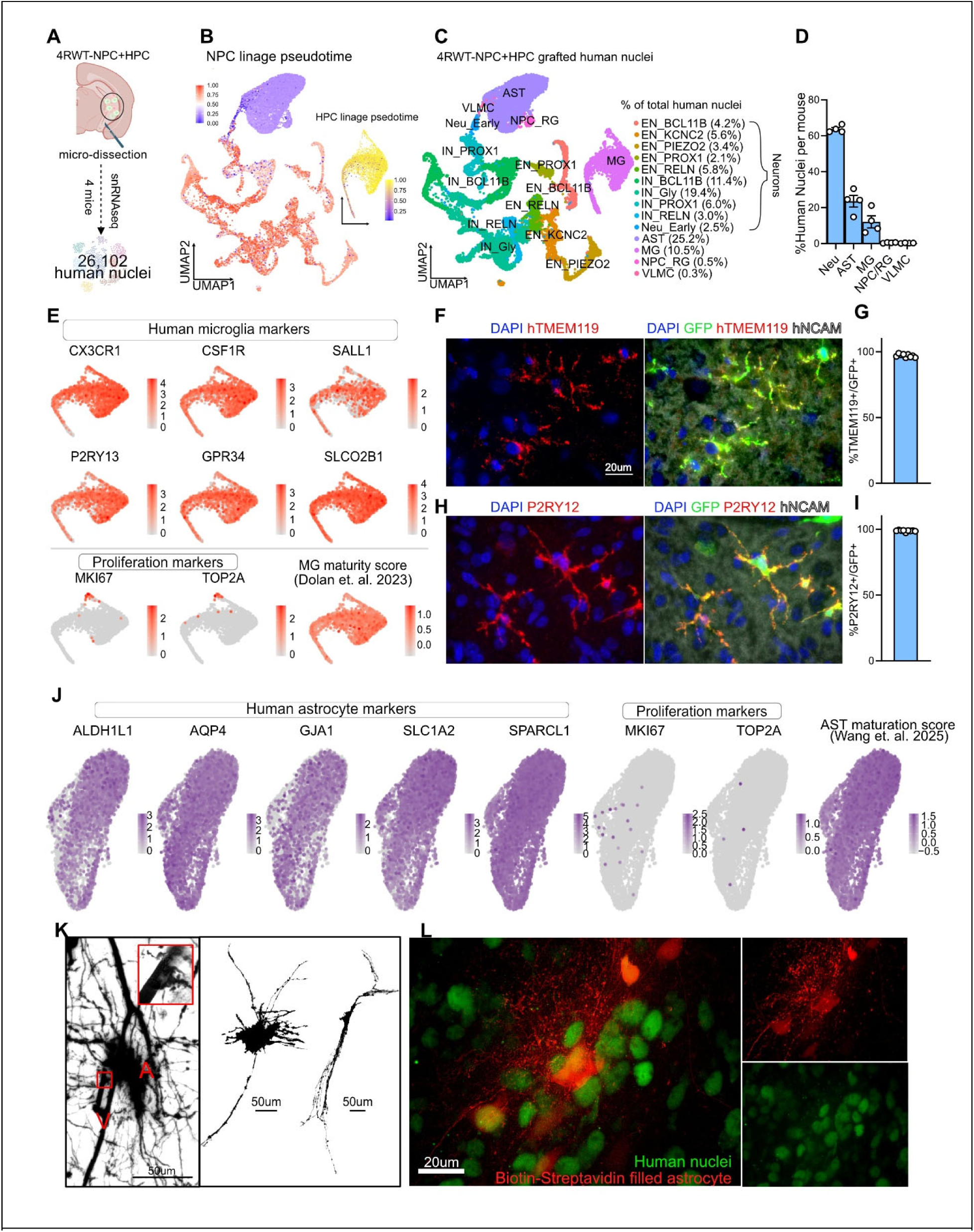
Single-nucleus profiling and characterization of microglia and astrocytes maturation in HuMiNAX. (**A**) Experimental schematic showing micro-dissection of 4R-WT NPC + HPC grafts from four mice followed by snRNA-seq profiling of 26,102 graft-derived human nuclei. (**B**) RNA velocity pseudotime analysis showing progressive differentiation of NPC-derived neural lineages and maturation trajectory of the HPC-derived microglial lineage. (**C,D**) UMAP visualization of graft-derived human nuclei colored by annotated cell type (C), with quantification of the proportional representation of major human cell classes per mouse (D). (**E**) Feature plots showing expression of canonical human microglial markers CX3CR1, CSF1R, SALL1, P2RY13, GPR34, and SLCO2B1, low expression of proliferation markers MKI67 and TOP2A, and enrichment of a human microglial maturity score (*27*) in graft-derived microglia. (**F,G**) Representative immunostaining showing expression of the homeostatic microglial marker TMEM119 in GFP⁺ human microglia within hNCAM⁺ human grafts (F), with quantification of the percentage of GFP⁺ cells expressing TMEM119 (G). N=9 mice. (**H,I**) Representative immunostaining showing expression of the homeostatic microglial marker P2RY12 in GFP⁺ human microglia within hNCAM⁺ human grafts (H), with quantification of the percentage of GFP⁺ cells expressing P2RY12 (I). N=9 mice. (**J**) Feature plots showing expression of human astrocyte identity markers ALDH1L1, AQP4, GJA1, SLC1A2, and SPARCL1, low expression of proliferation markers MKI67 and TOP2A, and enrichment of an astrocyte maturation score (*31*) in graft-derived astrocytes. (**K**) Golgi-Cox staining and representative reconstructions showing mature astrocyte-like morphology within human neural grafts, including astrocytic processes associated with a nearby vessel. A, astrocyte; V, vessel. (**L**) Three-dimensional reconstruction and post hoc visualization of a biocytin-filled human astrocyte counterstained with human nuclei, confirming the human origin and complex process morphology of graft-derived astrocytes. Labeling of nearby neurons is also observed, consistent with possible transfer of biocytin to adjacent cells through gap junctions.

Unsupervised clustering resolved major human cell populations, including excitatory and inhibitory neurons, astrocytes, and microglia (**Fig. 2C,D**). Neurons comprised ∼64% of human nuclei and included multiple excitatory and inhibitory subtypes marked by BCL11B, KCNC2, PIEZO2, PROX1, RELN, and glycinergic neuron signatures. Astrocytes and microglia accounted for ∼25% and ∼10% of nuclei, respectively, whereas NPC/radial glia-like and vascular leptomeningeal-like cells each represented <1%, indicating minimal persistence of progenitor populations four months after engraftment (**Fig. 2C,D**). This cellular composition closely matched histological analyses showing abundant human neurons, astrocytes, and microglia within HuMiNAX grafts (**Fig. 1D-J**). Notably, mouse microglia were absent from the snRNA-seq dataset, consistent with histological evidence of complete replacement of endogenous mouse microglia by engrafted human microglia within the grafted region (**Fig. 1K**).

We next examined human microglial identity and maturation markers. Graft-derived human microglia expressed CX3CR1 and CSF1R, together with CNS niche-associated genes including SALL1, P2RY13, GPR34, and SLCO2B1 (*25–27*), while showing minimal expression of MKI67 and TOP2A **(Fig. 2E)**. Broad expression of these markers and enrichment of a human microglial maturity score (*27*) indicate that grafted human microglia acquire features of a CNS conditioned maturation state in vivo. Consistent with this, GFP+ human microglia within hNCAM+ grafts expressed TMEM119 and P2RY12 by immunostaining, with nearly all GFP+ cells positive for these markers **(Fig. 2F-I)**. Together, these data support efficient engraftment and in vivo maturation of HPC-derived human microglia in HuMiNAX.

Graft-derived human astrocytes expressed ALDH1L1, AQP4, GJA1, SLC1A2, and SPARCL1(*28–30*), showed minimal proliferation marker expression, and were enriched for an astrocyte maturation signature (*31*)(**Fig. 2J**). These profiles indicate acquisition of mature astrocyte functions, including water homeostasis, gap junction coupling, glutamate uptake, and neuron-astrocyte interactions. Golgi-Cox staining further revealed astrocytes with elaborate processes and vascular-associated endfoot-like structures (**Fig. 2K**). Biocytin filling with HNA staining confirmed the human origin and complex morphology of graft-derived astrocytes, while biocytin transfer to neighboring neurons suggested astrocyte-neuron coupling within the graft (**Fig. 2L**).

Notably, 3R NPC + HPC grafts showed similar cellular maturation. snRNA-seq recovered 19,567 human nuclei from three mice and resolved comparable neuronal, astrocytic, microglial, NPC/radial glial, and VLMC populations (**fig. S4A,B**). Microglia and astrocytes derived from 3R NPC + HPC grafts expressed canonical lineage markers and were enriched for the corresponding maturation signatures (**fig. S4C,D**). These data demonstrate that HuMiNAX reproducibly generates diverse human neuronal subtypes, mature astrocytes, and CNS-conditioned microglia across NPC lines with distinct MAPT isoform backgrounds. This establishes HuMiNAX as a human neuroimmune platform suitable for modeling tauopathy-relevant cellular interactions in vivo.

## Tau propagation induces selective vulnerability of human neurons in HuMiNAX

Our previous studies established robust seeded tau aggregation in 4R iNeurons carrying an FTD-linked P301S mutation in culture (*14*). To model human neuroimmune responses to tau propagation in vivo, we co-engrafted G795A-GFP human HPCs with 4R-WT or 4R-P301S human NPCs, with or without human tau seeds, and analyzed grafts four months later (**Fig. 3A**). Human graft regions were defined by HNA staining, and pathological tau was quantified by chromogenic MC1 DAB staining to minimize autoflorenscence (*32*). Because MC1 recognizes misfolded endogenous tau but not recombinant tau seeds (*14, 33*), MC1 positivity indicates de novo tau misfolding and propagation.

**Fig. 3.**
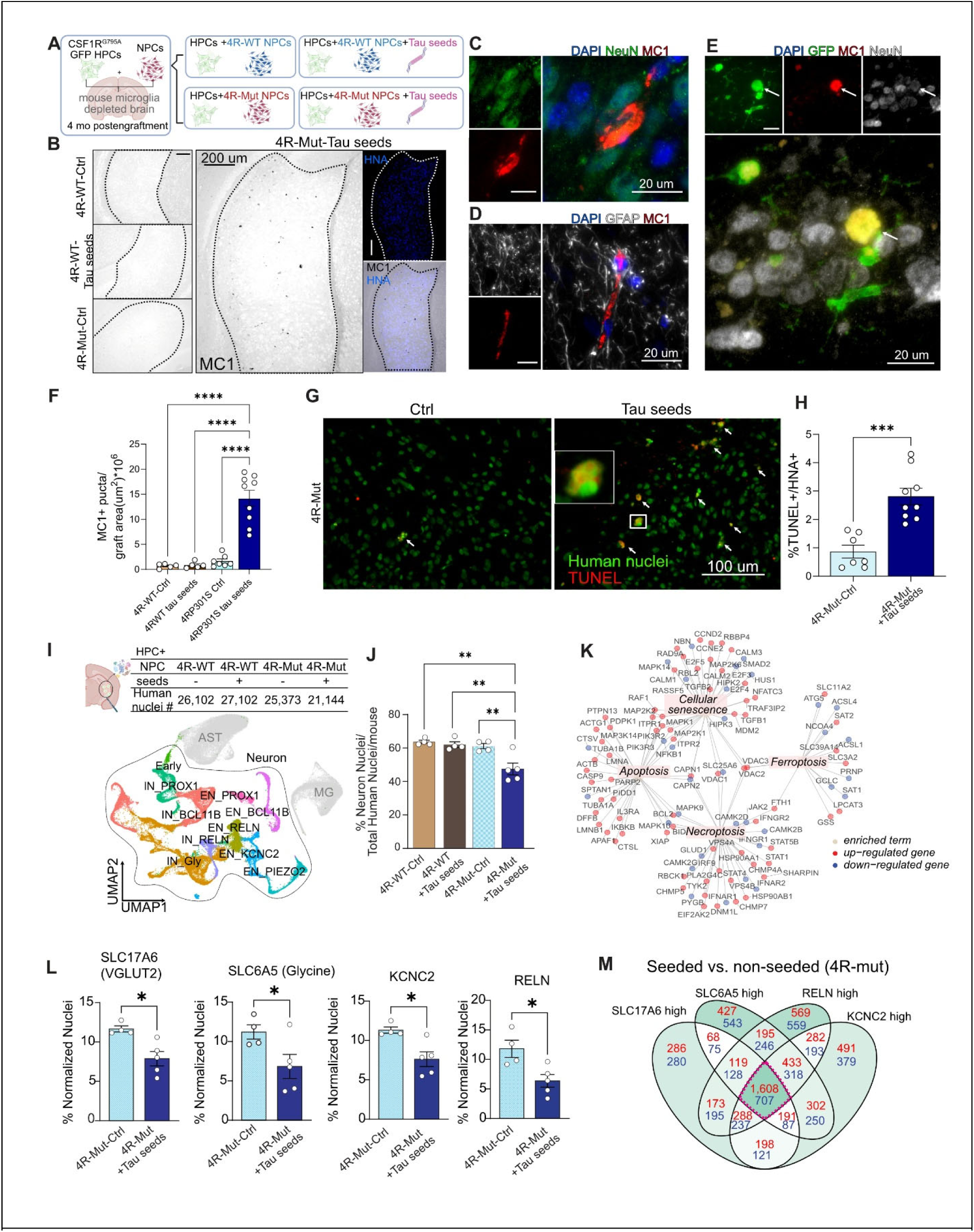
Tau seeding induces pathological tau accumulation, neuronal cell death, and selective neuronal vulnerability in HuMiNAX. (**A**) Experimental design. Human GFP-labeled hematopoietic progenitor-derived microglia were co-transplanted with 4R-WT or 4R-P301S (4R-Mut) neural progenitors into mouse microglia-depleted adult brains, with or without human tau seeds, and analyzed 4 months after grafting. (**B**) Representative MC1 immunostaining showing robust pathological tau accumulation in tau-seeded 4R-P301S ((4R-Mut) grafts, with minimal MC1 signal in unseeded 4R-WT, tau-seeded 4R-WT, or unseeded 4R-P301S (4R-Mut) grafts. Dashed outlines indicate graft regions, indicated by HNA+ area. (**C-D**) Immunofluorescence staining showing that MC1-positive pathological tau localizes predominantly to human NeuN-positive neurons, with limited association with GFAP-positive astrocytes. (**E**) Representative images showing GFP-positive human microglia surrounding or contacting MC1-positive human neurons. (**F**) Quantification of MC1-positive pathological tau burden across graft conditions, measured as MC1-positive puncta normalized to graft area. N=5-9 mice/group. (**G, H**) Representative images and quantification of TUNEL staining in HNA+ graft cells from unseeded and tau-seeded 4R-P301S (4R-Mut) grafts. Tau seeding increased TUNEL-positive human cells, indicating elevated graft cell death. N=7-9 mice/group. (**I**) Single-nucleus RNA sequencing of human graft nuclei across experimental conditions. UMAP visualization shows major human graft-derived cell populations and neuronal subtypes. Table indicates the number of recovered human nuclei per condition. (**J**) Quantification of the proportion of neuronal nuclei among total human nuclei per mouse, showing a selective reduction of human neurons in tau-seeded 4R-P301S grafts. (**K**) Term-gene graph of cell death-associated pathways and differentially expressed genes in tau-seeded versus unseeded 4R-P301S (4R-Mut) grafts, highlighting enrichment of cellular senescence, apoptosis, necroptosis, and ferroptosis-related programs. (**L**) Quantification of vulnerable neuronal populations in unseeded and tau-seeded 4R-P301S grafts. Tau seeding reduced SLC17A6-high, SLC6A5-high, KCNC2-high, and RELN-high neuronal populations. N=4-5 mice/group. (**M**) Venn diagram showing overlapping differentially expressed genes across vulnerable neuronal subpopulations in tau-seeded versus unseeded 4R-P301S (4R-Mut) grafts. Red numbers indicate upregulated genes and blue numbers indicate downregulated genes. Data are presented as mean ± SEM. Each dot represents one mouse. Statistical significance was determined by one-way ANOVA with post hoc multiple-comparison testing for multi-group comparisons and two-tailed unpaired t-test for two-group comparisons. *P < 0.05, **P < 0.01, ***P < 0.001, ****P < 0.0001.

Robust MC1 pathology was detected only in tau-seeded 4R-P301S grafts, whereas unseeded 4R-P301S, seeded 4R-WT, and unseeded 4R-WT grafts showed minimal signal **(Fig. 3B,F**), consistent with our in vitro findings (*14*). MC1 localized mainly to NeuN+ human neurons, with <1% detected in GFAP+ astrocytes (**Fig. 3C,D**). GFP+ human microglia frequently surrounded MC1+ neurons and contained NeuN- and MC1-positive material, suggesting uptake of tau-bearing neuronal debris (**Fig. 3E**). Tau propagation was accompanied by neurodegeneration. Tau-seeded 4R-P301S grafts showed increased TUNEL+ human nuclei compared with unseeded controls (**Fig. 3G,H**). This phenotype was reproduced in an independent KOLF 4R-P301S iPSC line, where tau seeding similarly induced MC1 pathology and TUNEL positivity (**fig. S5A-H**).

To define cellular responses to tau propagation, we performed snRNA-seq on high-quality human nuclei from 4R-WT and 4R-P301S grafts with or without tau seeding (**fig. S6A-N**). Using cell type labels defined in 4R-WT grafts as a reference, we annotated human nuclei across all conditions and detected similar major cell populations, including neurons, astrocytes, microglia, as well as less than 1% of NPC/radial glia-like cells and VLMCs. (**Fig. 3I; fig. S7A-D**). Tau-seeded 4R-P301S grafts showed reduced neuronal representation among total human nuclei, consistent with selective neuronal loss during tau propagation (**Fig. 3J**). Non-neuronal populations remained largely stable, although microglia showed a relative increase (**fig. S7E-H**). Differential expression and pathway analyses of tau-seeded versus unseeded 4R-P301S grafted human neurons revealed activation of senescence, apoptosis, necroptosis, and ferroptosis pathways (**Fig. 3K**), indicating a broad neurodegenerative transcriptional response.

We next asked whether tau-induced neuronal loss showed subtype selectivity. Tau-seeded 4R-P301S grafts showed preferential depletion of KCNC2-high, RELN-high, SLC6A5-high, and glycinergic neurons, whereas PROX1, PIEZO2, and BCL11B subtypes were relatively spared (**Fig. 3L; fig. S8A-L**). To define programs associated with vulnerable neurons, we analyzed shared differentially expressed genes (DEGs) across SLC17A6-high, SLC6A5-high, KCNC2-high, and RELN-high populations (**Fig. 3M**). Upregulated genes were enriched for protein catabolism, ubiquitin-proteasome function, translation, and mitochondrial respiration, whereas downregulated genes were enriched for synaptic organization, neurotransmission, vesicle localization, and membrane docking (**fig. S8M,N**). Thus, tau propagation induces neuronal stress and synaptic suppression programs, with selective vulnerability of defined human neuronal subtypes in vivo.

## Tau propagation reprograms HuMiNAX microglia

To define human microglial responses to tau pathology in vivo, we analyzed graft-derived microglia following tau seeding. Differential expression and pathway analyses revealed a shift toward an activated immune state, with upregulation of antigen presentation, Fcγ receptor-mediated phagocytosis, phagosome and lysosomal pathways, and inflammatory signaling programs, including chemokine and JAK-STAT signaling (**Fig. 4A**). In contrast, pathways associated with homeostatic and trophic functions, including neurotrophin, ErbB, MAPK, mTOR, FoxO, and lipid signaling, were reduced (**Fig. 4B**).

**Fig. 4.**
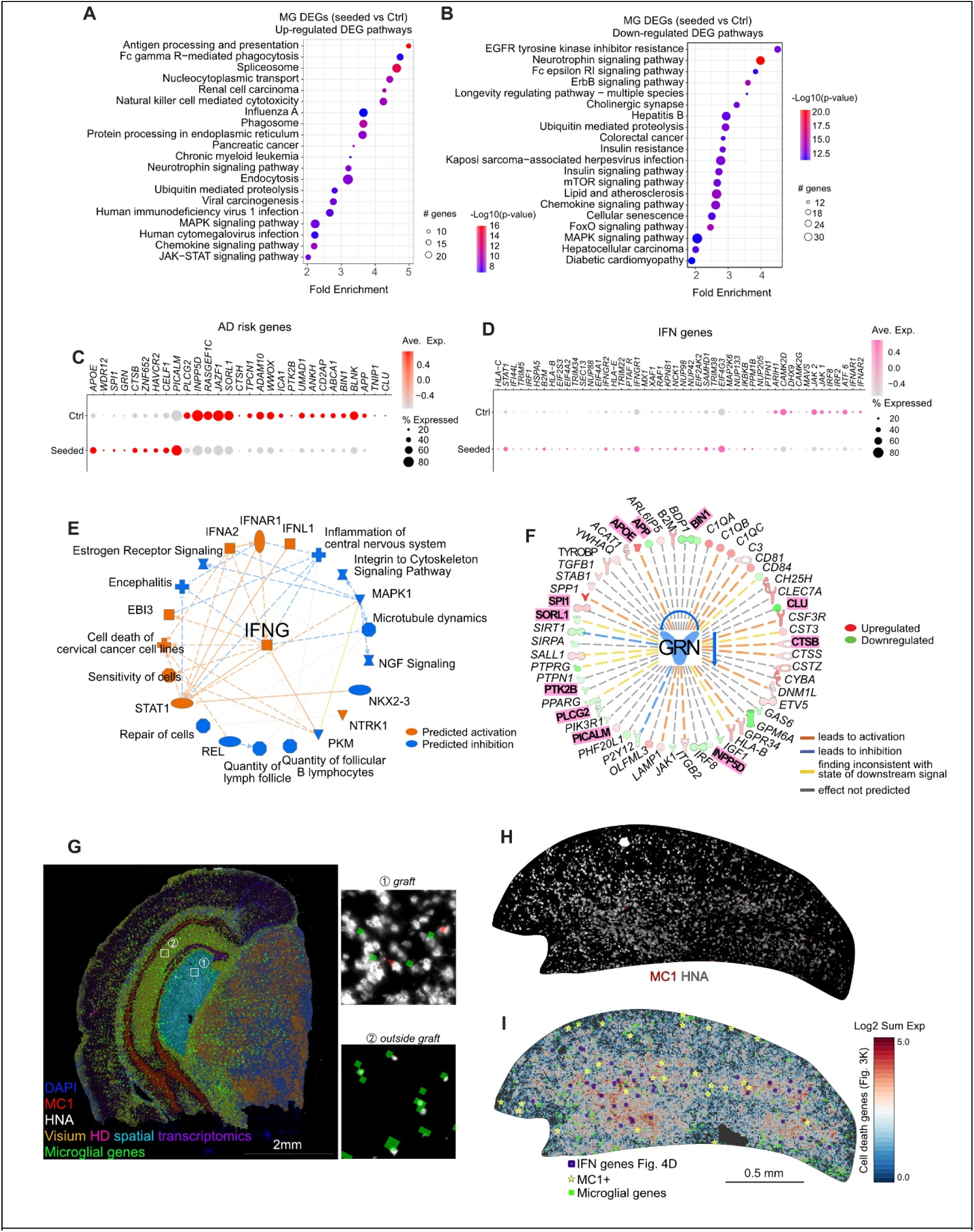
Tau seeding-induced tau aggregation elicit a human microglial interferon and disease-associated response in HuMiNAX. (**A-B**) PathfindR pathway enrichment analysis of upregulated (A) and downregulated (B) differentially expressed genes in graft-resident human microglia from tau seeded 4R-Mut grafts compared with unseeded 4R-Mut control grafts. (**C**) Dot plot showing expression of AD risk genes among microglia DEGs in control and tau seeded 4R-Mut grafts. Dot size indicates the percentage of microglia expressing each gene, and color indicates scaled average expression. (**D**) Dot plot showing expression of genes from the human Reactome interferon signaling gene set among microglia DEGs in control and tau seeded 4R-Mut grafts, revealing induction of an interferon-associated microglial program after tau seeding. (**E**) Ingenuity Pathway Analysis upstream regulator network identifying IFNG centered inflammatory signaling as a predicted regulator of tau seed induced microglial transcriptional changes. (**F**) Ingenuity Pathway Analysis of GRN related regulatory networks among microglial DEGs. Upregulated and downregulated genes are shown in red and green, respectively. Line color indicates predicted activation, inhibition, inconsistency with predicted downstream state, or no predicted effect. AD risk genes are bolded and highlighted in pink. (**G**) Representative Visium HD spatial transcriptomics section co-registered with DAPI, MC1, human nuclei antigen (HNA), and spatial expression of human microglial genes, showing broad distribution of graft associated human microglial signals within the human graft region. Insets show representative Visium HD spots inside and outside the graft. Microglial genes included *CX3CR1, SALL1, CSF1R, P2RY12, P2RY13, SLCO2B1, TMEM119, and SPI1*. (**H**) Spatial localization of MC1 pathology and HNA signal in a tau seeded HuMiNAX graft section used for spatial transcriptomic mapping. (**I**) Spatial mapping of the neuronal cell death associated gene program identified in Fig. 3K, overlaid with MC1 positive regions, interferon associated genes identified in Fig. 4D, and human microglial gene expression.

Tau-induced microglial changes also intersected with genetically implicated AD pathways, including increased *APOE* and *PICALM* expression and reduced *SORL1*, *INPP5D*, and *BIN1* expression (**Fig. 4C**) (*34*). Interferon-responsive genes were broadly induced (**Fig. 4D**), and upstream regulator analysis identified an IFNG-centered inflammatory network as a major driver of the microglial response (**Fig. 4E**). In parallel, IPA predicted suppression of a progranulin-associated regulatory network, with coordinated dysregulation of multiple *GRN*-linked genes (**Fig. 4F**). These findings suggest convergence of IFN activation and progranulin-related pathways on AD-relevant microglial programs.

To relate neurodegeneration to local neuroimmune signaling, we performed Visium HD spatial transcriptomics on graft-containing brain sections. Human microglial transcripts were broadly distributed throughout and around the graft (**Fig. 4G**). Overlaying neuronal cell-death signatures with MC1 pathology, IFN-associated microglial signatures, and human microglial gene expression revealed spatially restricted degeneration domains that aligned more closely with IFN-high regions than with MC1 pathology alone (**Fig. 4H,I, fig. S9A-D**). This pattern was reproduced in an independent sample (**fig. S9E-K**). This spatial dissociation suggests that neuronal vulnerability is linked more closely to local neuroimmune signaling and soluble tau than to visible tau inclusions.(*2, 3*).

## Progranulin-high human microglia in HuMiNAX restore resilience to tau toxicity

Loss-of-function mutations in *GRN* cause FTD and are linked to AD risk (*35, 36*). Although the 4R-P301S seeding model is not progranulin-deficient, prior work implicates PGRN in tau propagation and neurodegeneration (*35*), raising the possibility that enhancing PGRN expression in human microglia may confer protection across broader tauopathy contexts. Because tau seeding suppressed GRN-associated microglial networks, we next tested whether increasing PGRN specifically in human microglia could promote neuronal resilience during active tau propagation and evaluated engineered human microglia replacement as a potential strategy to modulate tau-driven neurodegeneration.

We inserted *GRN* open reading frame (ORF) downstream of the endogenous CX3CR1 locus in CSF1R^G795A^ human iPSCs to drive microglial PGRN overexpression (**Fig. 5A; fig. S10A-C**). Two independent lines, isogenic GRN-iso and non-isogenic GRN-2, showed normal karyotypes and were differentiated into HPCs for co-transplantation with 4R-P301S NPCs and tau seeds (**Fig. 5A; fig. S10D-F**).

**Fig. 5.**
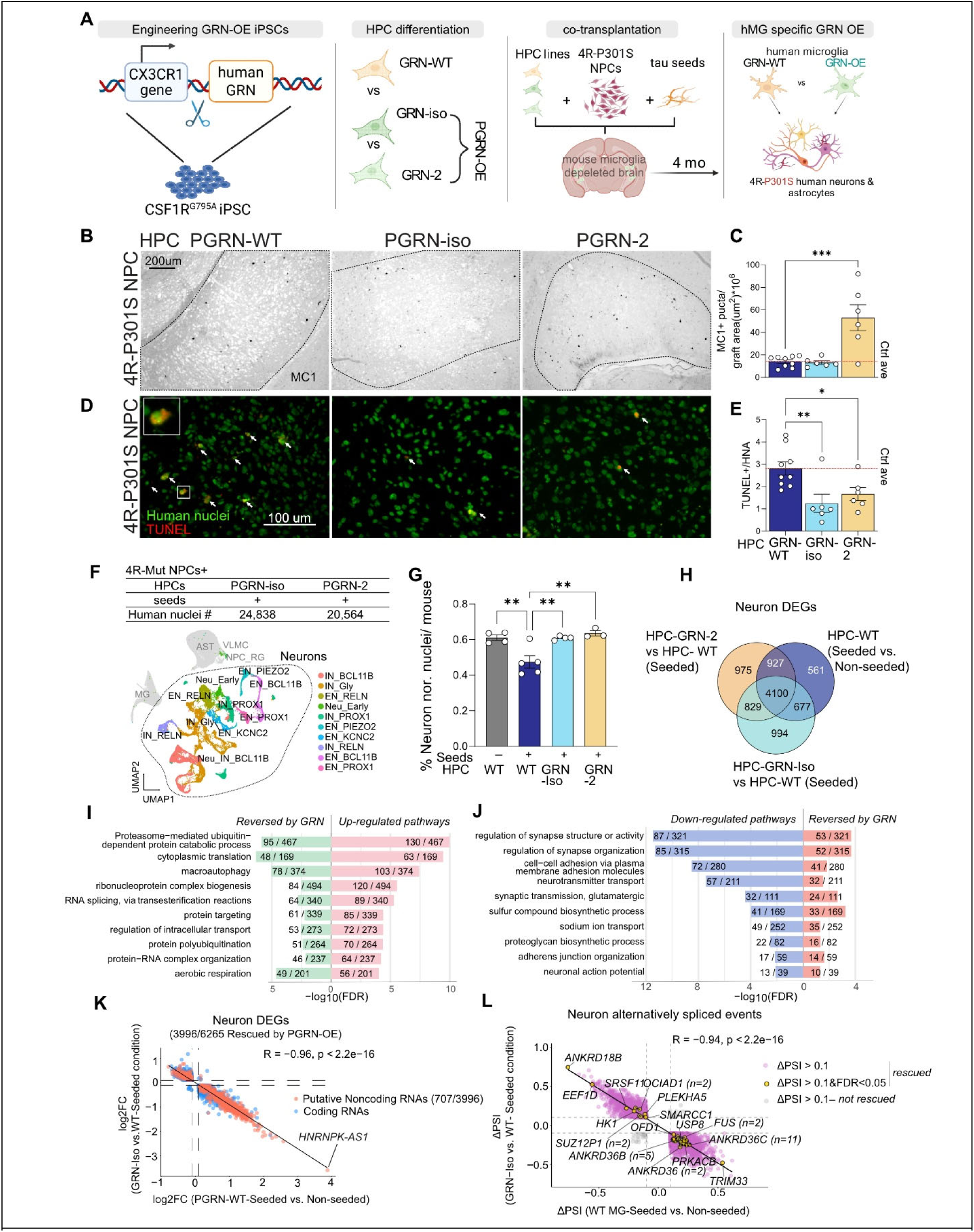
Microglial progranulin restores resilience to tau toxicity in HuMiNAX. (**A**) Experimental schematic. Human CSF1R^G795A^ iPSCs were engineered to overexpress human GRN from the endogenous CX3CR1 locus, differentiated into human microglia-like HPCs, and co-transplanted with 4R-P301S NPCs plus tau seeds into microglia-depleted mouse brains. Grafts were analyzed 4 months after transplantation to test the effect of microglia-specific progranulin overexpression on tau pathology and neuronal resilience. (**B-C**) Representative MC1 immunostaining and quantification of MC1-positive tau pathology in tau-seeded 4R-P301S HuMiNAX grafts containing WT, PGRN-iso, or PGRN-2 human microglia. The HPC-GRN-WT group is the same dataset shown in Fig. 3 and is replotted here as the seeded baseline reference for GRN-OE comparisons. Red line indicates the average value in non-seeded control grafts. HPC-GRN-WT, n = 9 mice; PGRN-iso, n = 6 mice; PGRN-2, n = 6 mice. One-way ANOVA followed by Dunnett’s multiple comparisons test comparing each condition with the seeded WT-HPC control column. **P < 0.01. (**D-E**) Representative TUNEL and human nuclear staining and quantification of TUNEL-positive human nuclei. Despite unchanged or increased MC1-positive tau pathology, grafts containing PGRN-overexpressing human microglia showed reduced human neuronal cell death compared with WT microglia. The HPC-GRN-WT group is the same dataset shown in Fig. 3 and is replotted here as the seeded baseline reference for GRN-OE comparisons. Red line indicates the average value in non-seeded control grafts. HPC-GRN-WT, n = 9 mice; PGRN-iso, n = 6 mice; PGRN-2, n = 6 mice. One-way ANOVA followed by Dunnett’s multiple comparisons test comparing each condition with the seeded WT-HPC control column. *P < 0.05, **P < 0.01. (**F**) UMAP visualization of human nuclei from tau-seeded 4R-P301S HuMiNAX grafts containing PGRN-iso or PGRN-2 microglia, re-integrated with the WT-HPC non-seeded and tau-seeded snRNA-seq datasets shown in Fig. 3I,J. Major human graft-derived populations and neuronal subtypes are annotated. (**G**) Quantification of the proportion of neurons among sequenced human nuclei showing tau seed-associated neuronal loss in WT-HPC grafts and preservation of neuronal representation in grafts containing PGRN-overexpressing microglia. WT-HPC non-seeded and WT-HPC tau-seeded conditions are replotted from Fig. 3I,J as reference groups. No-seed WT-HPC, n = 4 mice; seeded WT-HPC, n = 5 mice; PGRN-iso, n = 4 mice; PGRN-2, n = 3 mice. One-way ANOVA followed by Dunnett’s multiple comparisons test comparing each condition with the seeded WT-HPC control column. **P < 0.01. (**H**) Venn diagram showing overlap of neuronal differentially expressed genes across tau-seeded WT-HPC grafts and grafts containing PGRN-iso or PGRN-2 microglia. (**I-J**) Pathway analyses of neuronal DEGs showing tau-induced pathways that were reversed by GRN-OE microglia (I) and tau-suppressed neuronal pathways that were restored by GRN-OE microglia (J). (**K**) Correlation of neuronal log fold changes for consensus DEGs shared by both PGRN-OE microglial lines, comparing tau-seeded versus non-seeded WT-HPC grafts with PGRN-iso versus WT-HPC grafts under tau-seeded conditions. Directionally reversed genes indicate neuronal transcriptional programs induced by tau seeds and normalized by microglial GRN overexpression. Putative noncoding RNAs are highlighted, and HNRNPK-AS1 is indicated. (**L**) Single-nucleus isoform RNA sequencing (SnISOr-seq) analysis showing tau-associated alternative splicing changes in human neurons and their normalization by PGRN-iso microglia. Rescued splicing events were defined by opposite directional changes between tau-seeded versus non-seeded WT-HPC grafts and PGRN-iso versus WT-HPC grafts under tau-seeded conditions. Each point represents one alternatively spliced event. Gene labels indicate representative genes, with lowercase n denoting the number of splicing events assigned to that gene. SnISOr-seq was performed with N = 3 mice/group.

Four months later, both PGRN-OE lines showed increased PGRN in graft-resident human microglia by immunostaining and snRNA-seq (**fig. S10G,H**). MC1 staining showed that GRN-iso did not significantly alter tau pathology, whereas non-isogenic GRN-2 increased MC1 burden relative to WT-HPC controls (**Fig. 5B,C**). This difference may reflect donor-specific microglial genetic background that influences the clearance of MC1-positive tau pathology. However, despite these divergent effects on MC1 pathology, both PGRN-OE lines reduced TUNEL+ human nuclei (**Fig. 5D,E**), indicating protection from tau-associated cell death independent of visible tau burden.

We next performed snRNA-seq on microdissected PGRN-OE grafts and integrated these datasets with 4R-P301S control and 4R-P301S tau-seeded conditions, yielding high-quality human nuclei for downstream analyses (**fig. S6A-N**). snRNA-seq confirmed preservation of human neurons in PGRN-OE grafts. Tau seeding reduced neuronal representation in WT-HPC grafts, whereas grafts containing GRN-iso or non-isogenic GRN-2 microglia maintained neuronal proportions (**Fig. 5F,G; fig. S10I-K**). Thus, microglial PGRN overexpression reproducibly protects human neurons from tau-associated degeneration in vivo. We next defined transcriptional DEGs shared by the two neuroprotective PGRN-OE lines. GRN-iso and non-isogenic GRN-2 induced overlapping changes across microglia, astrocytes, and neurons under tau-seeded conditions (**Fig. 5H; fig. S11A-C**). In microglia and astrocytes, PGRN-OE broadly opposed tau-induced transcriptional changes and restored pathways related to endolysosomal transport, protein homeostasis, cytoskeletal organization, synapse organization, and lipid modification (**fig. S12A-H**). These findings indicate that microglial progranulin may promote neuronal resilience through both intrinsic and non-cell-autonomous mechanisms.

## Rescue of neuronal transcriptional program by progranulin-high human microglia

We then defined the neuronal transcriptional programs associated with PGRN-mediated resilience. In WT-HPC grafts, tau seeding induced neuronal pathways related to proteasome-mediated protein catabolism, macroautophagy, ribonucleoprotein complex biogenesis, RNA splicing, protein targeting, and aerobic respiration. These tau-induced programs were reversed by both GRN-iso and non-isogenic GRN-2 microglia **(Fig. 5I)**. Conversely, neuronal pathways suppressed by tau seeding, including synapse organization, neurotransmission, cell adhesion, glutamatergic signaling, sodium ion transport, and neuronal action potential pathways, were restored by PGRN-OE microglia **(Fig. 5J)**. This result suggests that microglial progranulin preserved neuronal transcriptional programs linked to synaptic identity and function while reducing stress-associated RNA and protein homeostasis responses.

To identify a reproducible neuronal resilience signature, we compared tau-induced neuronal DEGs in WT-HPC grafts with the consensus DEGs reversed by both PGRN-OE microglial lines. Overall, 3,996 of 6,265 tau-associated neuronal DEGs were rescued by PGRN-OE microglia, and the direction of PGRN-mediated change was strongly anti-correlated with the tau-seed effect (R = −0.96, **Fig. 5K**). Notably, the rescued neuronal gene set included a large fraction of putative noncoding RNAs. Among these, HNRNPK-AS1 emerged as one of the most strongly tau-induced and PGRN-rescued transcripts **(Fig. 5K)**. Because HNRNPK-AS1 is antisense to HNRNPK, an RNA-binding protein implicated in RNA toxicity, DNA damage responses, and cryptic splicing in ALS and FTLD (*37–39*), we reasoned that microglial PGRN may influence neuronal vulnerability through RNA regulatory mechanisms.

Consistent with this possibility, long-read single-nucleus isoform RNA sequencing (SnISOr-seq) identified tau-associated alternative splicing events in human neurons that were normalized by GRN-iso microglia **(Fig. 5L)**. These rescued splicing changes included events in genes linked to RNA processing, cytoskeletal regulation, and neuronal function. Thus, changing PGRN expression only in human microglia was sufficient to reshape neuronal transcriptomic and splicing programs, indicating that microglial state can non-cell-autonomously regulate RNA homeostasis in human neurons. Together, these data show that microglial progranulin restores neuronal resilience in tau-seeded HuMiNAX grafts by suppressing neurodegeneration and normalizing RNA regulatory, splicing, synaptic, and stress-response programs.

## A microglia-regulated neuronal lncRNA axis drives neurodegeneration

Our data revealed neuronal HNRNPK-AS1 as one of the strongest tau-induced transcripts reversed by PGRN-OE microglia. Because HNRNPK-AS1 is antisense to HNRNPK, an RNA-binding protein implicated in RNA processing, splicing, FTLD, and ALS (*37–39*), we tested whether HNRNPK is required for neuronal survival. HNRNPK knockdown in 4R-P301S human neurons reduced viability, induced RNA metabolic, proteostatic, mitochondrial, and cell-death programs, and suppressed synaptic, membrane-potential, neurotransmission, and vesicle-cycling pathways (**fig. S13A-G**). It also caused widespread splicing changes in genes linked to neuronal projections, excitability, autophagy, mitochondria, synapses, and vesicle organization (**fig. S13H,I**), consistent with previous reports (*37–39*).

Compared with HNRNPK, HNRNPK-AS1 is poorly characterized and not conserved in rodents (*11*), making it a particularly attractive human-specific candidate that cannot be readily modeled in conventional mouse systems. To test how microglial PGRN signaling may regulate HNRNPK-AS1, we scanned its proximal promoter using FIMO and the JASPAR2026 CORE vertebrate motif database (*40, 41*). This identified candidate transcription factors whose expression was altered by tau seeding and normalized by microglial PGRN-OE (**Fig. 6A,B**). Tau-induced and PGRN-suppressed factors included STAT1 and STAT5B, linking HNRNPK-AS1 regulation to interferon signaling, as well as NR2F1, MEIS3, EBF1, ESR2, and ZBTB11, implicating altered neuronal identity and stress programs (*24, 42–45*). Conversely, tau-suppressed and PGRN-restored factors included KLF7, KLF6, and ZNF841, consistent with recovery of neuronal maintenance programs (*46, 47*). Thus, HNRNPK-AS1 may define a tau-responsive neuronal state driven by microglial inflammatory and IFN signaling, with diminished resilience.

**Fig. 6.**
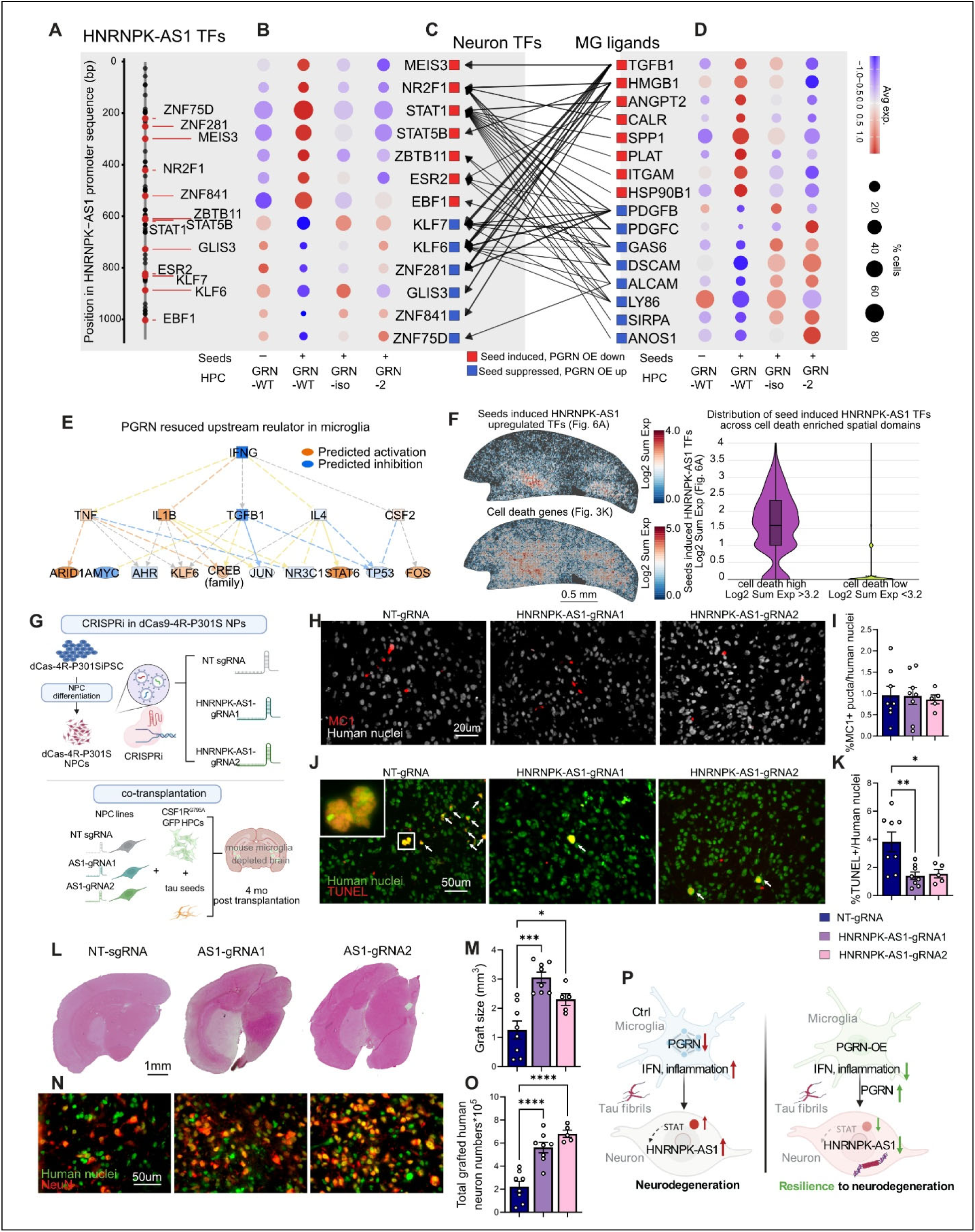
Microglial progranulin suppresses an HNRNPK-AS1 associated neuronal degeneration program and promotes resilience to tau toxicity. (**A**) FIMO predicted transcription factor (TF) binding motifs in the HNRNPK-AS1 promoter using the MEME Suite and JASPAR2026 CORE vertebrate nonredundant position frequency matrix database. Labeled motifs indicate top TFs associated with microglia regulated neuronal programs. (**B**) Dot plot showing expression of HNRNPK-AS1 promoter associated transcription factors in human neurons across control, tau seeded, and microglial PGRN OE conditions. (**C**) NicheNet mapping of neuronal HNRNPK-AS1 TFs to PGRN rescued microglial ligands. Red and blue side bars indicate seed induced, PGRN OE downregulated TFs or ligands, and seed suppressed, PGRN OE upregulated TFs or ligands, respectively. Connecting lines indicate predicted ligand to TF links. (**D**) Dot plot showing expression of top microglial ligands across control, tau seeded, and microglial PGRN OE conditions. (**E**) IPA predicted upstream regulator network from PGRN rescued microglial genes. Orange indicates predicted activation and blue indicates predicted inhibition. (**F**) Spatial transcriptomic maps of the seed induced HNRNPK-AS1 associated TF program and cell death related gene program, with violin plot of spatial expression distribution across cell death high and cell death low spatial domains. (**G**) Schematic of CRISPR interference strategy for HNRNPK-AS1 knockdown in dCas9 expressing 4R P301S human neural progenitor cells before co transplantation with human microglial precursors and tau seeds. (**H, I**) Representative images and quantification of MC1 positive pathological tau puncta normalized to human nuclei. (**J, K**) Representative images and quantification of TUNEL positive human nuclei. (**L, M**) Eosin staining and quantification of graft size. (**N, O**) Representative human nuclei and NeuN staining with quantification of total grafted human neurons. (**P**) Working model of microglial PGRN mediated regulation of inflammatory signaling, HNRNPK-AS1 associated neuronal transcriptional programs, and neuronal resilience against tau toxicity in tau seeded HuMiNAX grafts. Data are shown as mean ± SEM. Each dot represents one mouse. N = 5 to 8 mice per group. Statistical significance in I, K, M, and O was determined by ordinary one-way ANOVA followed by Tukey’s multiple comparisons test. *P < 0.05, **P < 0.01, ***P < 0.001, ****P < 0.0001.

To further test this hypothesis, we next examined HNRNPK-AS1 associated neuronal transcription factors as candidate downstream targets in the NicheNet analysis of microglia to neuron signaling (*48, 49*). The predicted microglial ligands rescued by PGRN-OE converged on damage associated inflammatory cues and phagocytic or homeostatic signaling pathways **(Fig. 6C,D)**. Among these, SPP1 and HMGB1 provided inflammatory damage response anchors (*50, 51*), whereas TGFB1, GAS6, and SIRPA pointed to a second axis involving microglial homeostasis, phagocytic regulation, and neuron microglia interaction (*52–54*). Therefore, the NicheNet analysis supported a model in which microglial PGRN-OE remodels inflammatory and phagocytic signaling programs that converge on neuronal transcription factors predicted to regulate HNRNPK-AS1. Consistent with STAT1 being identified as a seed-induced, PGRNOE-suppressed candidate regulator of the HNRNPK-AS1 promoter, IPA predicted reduced IFNG signaling among PGRN rescued microglial genes **(Fig. 6E)**. These findings suggest that PGRN-OE dampens microglial IFN signaling and inflammatory programs, thereby attenuating neuronal transcriptional activation, including STAT1-associated responses that may contribute to HNRNPK-AS1 elevation. Spatial transcriptomic mapping further showed that the seed induced HNRNPK-AS1 associated transcription factor program overlapped with spatial domains enriched for cell death related genes **(Fig. 6F)**. Quantification across spatial domains showed higher expression of this transcription factor program in cell death high regions than in cell death low regions, linking the HNRNPK-AS1 regulatory axis to neurodegeneration associated tissue domains in vivo.

We then directly tested whether HNRNPK-AS1 contributes to neuronal vulnerability in HuMiNAX. Using CRISPR interference, we targeted HNRNPK-AS1 in dCas9 expressing 4R-P301S human neural progenitor cells before co transplantation with human microglial precursors and tau seeds **(Fig. 6G)**. Four months after transplantation, two independent HNRNPK-AS1 guide RNAs reduced nuclear HNRNPK-AS1 signal in grafted human cells, as confirmed by RNAscope in situ hybridization **(fig. S14)**. HNRNPK-AS1 knockdown did not significantly alter MC1 positive pathological tau burden **(Fig. 6H,I)**. In contrast, both HNRNPK-AS1 guide RNAs reduced TUNEL positive human nuclei compared with non-targeting controls **(Fig. 6J,K)**. HNRNPK-AS1 knockdown also increased overall graft size **(Fig. 6L,M)** and increased the total number of grafted human neurons **(Fig. 6N,O)**. These findings suggest that HNRNPK-AS1 suppression enhances neuronal resilience downstream of tau accumulation without reducing detectable MC1 positive tau pathology.

Together, these results support a model in which tau seeding activates microglial interferon and inflammatory signaling and induces an HNRNPK-AS1 associated neuronal vulnerability program. Microglial PGRN OE suppresses this inflammatory and HNRNPK-AS1 associated neuronal state, while direct CRISPRi mediated HNRNPK-AS1 knockdown is sufficient to reduce neuronal cell death and preserve grafted human neurons without lowering MC1 positive tau burden **(Fig. 6P)**. Thus, HNRNPK-AS1 defines a human relevant, microglia regulated neuronal lncRNA axis that can be functionally targeted in HuMiNAX to promote neuronal resilience in tau seeded human neural grafts.

## Discussion

Modeling human tauopathies remains challenging because tau propagation, neuroimmune responses, and neuronal loss are highly context dependent and can diverge across species (*55*). Although mouse models have provided essential mechanistic insights, human brain cells show distinct transcriptional responses to tau pathology, and many lncRNA networks are poorly conserved between humans and mice (*11*). Here, we establish HuMiNAX, an in vivo humanized neuroimmune platform that integrates human neurons, astrocytes, and microglia while replacing endogenous murine microglia. This perturbable iPSC-based system enables cell-type-resolved profiling and functional testing of human microglia-neuron interactions during tau propagation.

HuMiNAX extends prior transplantation approaches by combining scalable human NPC grafting with human microglia replacement in the adult brain models (*18–22*). Earlier dissociated human NSC or NPC transplantation studies established that human progenitors can survive in vivo, but have often been limited by incomplete terminal differentiation, ectopic growth, restricted electrophysiological maturation, or limited long term neuronal survival (*56–58*). Human organoid grafts have shown that the brain environment can promote vascularization, neuronal maturation, axonal projection, and circuit integration (*31, 59–61*), but organoid transplantation can be difficult to standardize and adapt for coordinated maturation of neurons, astrocytes, and microglia. HuMiNAX uses defined, cryobankable, genetically engineerable iPSC-derived NPCs (*18*) and HPCs, enabling reproducible cohort design, tau seeding, single-nucleus profiling, and cell-type-specific perturbation. Although the grafts remain spatially restricted, this feature facilitates microdissection and selective analysis of human graft compartments. The resulting grafts contained diverse neuronal subtypes, mature astrocytes, CNS-conditioned human microglia, dendritic spine development, and electrophysiologically active neurons, establishing a controlled in vivo platform to study human neuroimmune mechanisms.

Tau propagation shifted graft-derived human microglia toward antigen-presentation, phagocytic, lysosomal, chemokine, JAK-STAT, and interferon programs, while reducing trophic and homeostatic pathways. This disease-associated state may have adaptive or maladaptive consequences depending on timing, context, and neuronal vulnerability (*62*). In HuMiNAX, the spatial association between IFN-rich domains and neurodegeneration suggests that sustained inflammatory microglial signaling becomes harmful during tau propagation (*24, 63*).

The protective effect of xenografted progranulin-high human microglia places PGRN within this neuroimmune resilience framework. GRN haploinsufficiency causes FTD and has been linked to AD risk (*35, 36, 64*). Notably, neuronal protection occurred despite divergent effects on MC1-positive pathology across two GRN-OE lines. This indicates that microglial PGRN does not simply block tau seeding or clear visible tau deposits. Instead, PGRN appears to buffer downstream toxic responses by restoring neuronal transcriptional and splicing programs, preserving synaptic pathways, and dampening glial stress states. These findings support therapeutic strategies that elevate PGRN to enhance neuronal resilience rather than only reduce tau burden (63).

An unexpected implication is that microglia can regulate neuronal RNA homeostasis in vivo. Altering GRN only in human microglia reshaped neuronal programs related to RNA processing, splicing, synaptic function, stress responses, and neuronal identity. HNRNPK-AS1 emerged from this response as a human-relevant lncRNA at the intersection of tau toxicity and microglia-mediated resilience. Its induction in vulnerable neurons, suppression by progranulin-high microglia, and weak conservation in rodents highlight the importance of humanized systems for discovering regulatory pathways that may be invisible in conventional mouse models (*11*).

Functional perturbation of HNRNPK-AS1 further separated neuronal resilience from tau deposition. CRISPRi-mediated knockdown reduced neuronal cell death and preserved graft-derived human neurons without lowering MC1-positive tau burden. This places HNRNPK-AS1 downstream of tau accumulation and suggests that tau-induced neuronal death can be modified by targeting RNA regulatory states rather than tau pathology itself. Because HNRNPK-AS1 is antisense to HNRNPK, an RNA-binding protein involved in RNA processing, alternative splicing, DNA damage responses, and neurodegenerative disease mechanisms (*37–39*), this locus may link tau-induced neuroimmune stress to neuronal RNA dysregulation.

Several limitations remain. HuMiNAX currently lacks human oligodendrocytes, which will be important for modeling myelination, white matter vulnerability, and glial interactions relevant to broader neurodegenerative disease. Efficient human oligodendrocyte engraftment may require more permissive myelination niches or demyelination paradigms (*65, 66*). Stable xenograft survival also requires immune-permissive hosts and depletion of endogenous mouse microglia, limiting analysis of adaptive immunity and intact peripheral immune responses. In addition, the predicted pathway linking microglial inflammatory ligands, neuronal STAT-linked transcriptional programs, and HNRNPK-AS1 induction requires direct mechanistic testing. Finally, although both male and female hosts were used, the human donor lines were male; future studies should systematically test sex-related effects in human graft-derived cells.

Overall, this study establishes HuMiNAX as an in vivo humanized platform for modeling tau propagation, dissecting microglia specific regulation of neuronal vulnerability, and functionally testing human specific disease associated regulatory elements. Our findings identify microglial PGRN as a non-cell autonomous resilience factor and HNRNPK-AS1 as a neuronal lncRNA vulnerability axis downstream of tau induced neuroimmune stress. Together, these results support a model in which neuronal loss can be uncoupled from visible tau burden and instead shaped by local microglial signaling, neuronal RNA regulatory states, and human specific lncRNA programs.

## Materials and methods

### iPSC maintenance and differentiation

Human iPSCs were maintained and described in previous studies(*14, 67*), which harbor a tet-on NGN2 construct but doxycycline was not added to differentiation steps in generating neural progenitor cells (NPCs) or hematopoietic progenitor cells (HPC) used for transplantation. Human iPSCs were maintained on a Matrigel coated plates supplemented with mTeSR Plus medium (STEMCELL, #05825) and passaged using ReLeSR (STEMCELL, # 05872). The 4R-WT, 4R-

P301S, and dCas9-4R-P301S iPSC lines were previously characterized, and the KOLF2.1J line (The Jackson Laboratory, JIPSC001000) and KOLF 4R-P301S iPSC line were engineered as described in our previous study (*14*).

To generate a stable iPSC line that overexpresses progranulin (PGRN) upon microglial differentiation, we constructed a CX3CR1 targeted donor plasmid in which a T2A GRN cDNA polyA cassette and an EF1α puromycin GFP selection cassette were flanked by the left and right homology arms of the human CX3CR1 gene, enabling in frame insertion of T2A GRN immediately upstream of the CX3CR1 stop codon. The CX3CR1 homology arms were synthesized based on CX3CR1 genomic sequences and cloned into the donor plasmid to minimize the impact of potential single nucleotide polymorphisms, and an sgRNA targeting the 3′ end of CX3CR1 was selected using the IDT CRISPR design tool. Cas9 ribonucleoprotein complexes and donor DNA were electroporated into iPSCs using the Lonza Nucleofector system, followed by puromycin selection for 5 days and establishment of single cell derived clones by plating individual cells into 96 well plates. Correct on target integration was confirmed by junction PCR using primers located outside the homology arms in the genome and within the inserted cassette, and PCR products from positive clones were validated by Sanger sequencing; clones were additionally screened to exclude nonspecific or random integration events, and homozygous knock in clones were identified by PCR genotyping and expanded. For WTC11 derived CSF1R G795A iPSC lines, which already carried the G795A variant, PGRN knock in was introduced directly into the pre-existing CSF1R G795A background to generate isogenic PGRN OE lines. In parallel, a male WiCell iPSC line (UCSD075i-1-6) was first engineered with the CX3CR1 T2A GRN knock in cassette, and the CSF1R G795A variant was subsequently introduced following a published protocol (*20*) using crRNA CSF1R G795 (5′ ATGGTCATGTGGCCAAGATT 3′) and an ssODN donor (5′ CAGTGCATCCACCGGGACGTGGCAGCGCGTAACGTGCTGTTGACCAATGGTCATGTGGCCAAGATTGCCGACTTCGGGCTGGCTAGGGACATCATGAATGACTCCAACTACATTGTCAAGGGCAATGTAAGTGCTGGGAGGGCTTGGGCC 3′). All edited iPSC clones were further validated by karyotyping and assessment of pluripotency marker expression prior to downstream differentiation.

NPCs were generated from iPSC lines as previously described (*18, 23, 68*). HPC were differentiated from iPSCs using the STEMdiff™ Hematopoietic Kit (STEMCELL, #05310) following manufacturer’s instructions. HPCs used for transplantation were freshly harvested on days 11 and 13 of differentiation.

NPCs were differentiated into doxycycline induced NGN2 cortical neuron similar to previously described (*14*). Briefly, NPCs were dissociated with Accutase and seeded onto PDL coated plates at the density of 0.4 million cells/ml in NPC basal medium (equal mix of DMEM-F12 and Neurobasal A with 0.5X B27 without vitamin A, 0.5X N2, 1% penicillin streptomycin with glutamine) supplemented with 1x non-essential amino acids (Gibco, 11140050), 10ng/ml BDNF (PeproTech, 450-02), 10ng/ml of NT3 (PeproTech, 450-03), 1µg/ml of mouse laminin (Trevigen, 3446-005-01), and 2 ug/ml doxycycline (Sigma, D9891). Half medium was changed every 3 days, and after a week, neurons were maintained in the same medium without doxycycline.

### PLX treatment

PLX5622 FA was obtained from WuXi and converted to PLX5622-FA-CC as a dry powder prior to chow preparation, as described previously (*69*). Briefly, poloxamer 407 (20.0% wt/wt) and crospovidone XL10 (1.5% wt/wt) were combined in a mortar and premixed until uniform. PLX5622 FA (78.0% wt/wt) was then incorporated in portions, allowing 2 to 3 min per addition, followed by gentle grinding with a pestle in a circular motion for 5 min after each addition. The blend was mixed for an additional 2 to 3 min to disrupt any visible agglomerates. Magnesium stearate (0.5% wt/wt) was added last and mixed until a homogeneous powder was obtained. This PLX5622-FA-CC powder mix was then incorporated into mouse chow by Research Diets, Inc. to achieve a final PLX5622 concentration of 1500 mg/kg diet. Mice were placed on the PLX5622 diet 2 weeks prior to human cell transplantation and maintained on the diet for an additional 1.5 months after stereotaxic surgery.

### Transplantation in mice

All animal experiments were conducted in the Research Animal Resources and Compliance facility (RARC) at Weill Cornell Medicine, and all procedures were performed in accordance with RARC standards for animal care under IACUC approved protocol: 2018-0043. Xenograft competent humanized immunodeficient mice (hCSF1^KI/KI^-Rag2^-/-^-Il2rg^y/-^) were purchased from Jackson Laboratory (Strain #:017708). Mice were housed in a barrier facility under a 12-hour light/12-hour dark cycle with controlled temperature conditions. Per institutional protocol, xenotransplanted animals were transferred to a dedicated biohazard xenograft housing room following surgery and maintained there until tissue harvest.

NPCs were fed with fresh medium at least one hour before dissociation with Accutase to generate a single-cell suspension and then washed twice with PBS. HPCs were harvested by gentle pipetting to collect floating cells in HPC medium and then washed twice with PBS. Live NPC and HPC counts were determined using a Countess II automated cell counter (Invitrogen Countess II) with trypan blue counterstaining. Cells were then resuspended in sterile PBS at a density of 35,000 cells/µl per cell type and kept on ice until stereotaxic surgery.

For tau seeded conditions, tau seeds (tau (K18) P301L mutant pre-formed fibrils; StressMarq Biosciences, SPR-330) were sonicated in an ice bath for a total of 10 min using alternating 30s on and 30 s off pulses at 40% amplitude with a water bath sonicator (EpiSonic 2000, EpigenTek). Sonicated tau seeds were then added to the HPC plus NPC cell suspension to a final concentration of 0.5 mg/ml.

The cell suspension was stereotaxically injected into the bilateral hemispheres (coordinates: x ±2.9, y −3.2, z −3.0) of 2 to 3 months old young adult mice that were deeply anesthetized with 2% isoflurane and secured in a stereotaxic frame (Kopf Instrument), flowing stereotaxic procedures approved by RARC. Three microliters were injected into each side of the brain hemisphere at the speed of 400nl/min.

### Immunohistochemistry of human grafts and imaging

Mice were euthanized by intraperitoneal (IP) injection of pentobarbital solution (Euthasol) and perfused transcardially with cold PBS for 5 min. After complete perfusion, brains were dissected and one hemibrain was fixed in 4% paraformaldehyde overnight at 4 °C, then transferred to 15% sucrose in PBS containing 0.05% sodium azide and stored at 4 °C overnight or until sectioning. On the day of cryostat sectioning, brains were snap-frozen in dry-ice pre-cooled isopentane (2-methylbutane; Sigma, M32631) and sectioned at 10 µm thickness using a cryostat (Leica). Serial sections were mounted onto charged glass slides until human grafts were no longer visible and stored at −20 °C until use.

Eosin staining was performed using the Hematoxylin and Eosin (H&E) Staining Kit (Abcam, ab245880). Because human and mouse cells differ only in cytosolic content, the hematoxylin step was omitted. Sections were hydrated in distilled water for 5 min, dehydrated in absolute ethanol for 1 min, submerged in Eosin Y solution for 3 min, rinsed with absolute ethanol, and mounted with Permount mounting medium (Fisher, SP15-500). Unbiased estimates of transplant volume were obtained in ImageJ by analyzing stained serial 10 μm brain sections. For quantification, every tenth 10 μm section spanning the graft was systematically sampled and analyzed.

Immunofluorescent staining was performed as previously described(*18*). Briefly, frozen sections were thawed to room temperature, washed three times in PBS, and blocked for 1 h in 5% normal donkey serum (NDS) with 0.2% Triton X-100 in PBST, followed by overnight incubation with primary antibodies at 4 °C. Slides were washed three times in PBS the following day, incubated with Alexa Fluor-conjugated secondary antibodies (Invitrogen) for 1 h at room temperature, washed three additional times in PBS, and mounted in antifade mounting medium containing DAPI (Vector Laboratories, H-1500-10). Fluorescent images were acquired using a Keyence fluorescence microscope (bz-x710). Quantification was performed in QuPath using a supervised pixel classifier. Multiple representative images from each experimental condition were used to train the classifier, and a single finalized classifier was then applied uniformly across all samples to quantify staining in an unbiased manner. All images were processed using identical detection and measurement parameters across conditions.

MC1 DAB staining. MC1 chromogenic detection was performed following immunofluorescent staining, immediately after secondary antibody incubation. Sections were quenched in 3% H₂O₂ for 10 min, then sequentially blocked with avidin and biotin solutions for 15 min each (Vector Laboratories, SP-2001). The M.O.M. Immunodetection Kit (Vector Laboratories, BMK-2202) was applied per the manufacturer’s instructions: sections were incubated with MC1 primary antibody (1:1000) for 1 h at room temperature, followed by biotinylated anti-mouse IgG secondary antibody for 30 min. After three PBS washes, VECTASTAIN Elite ABC Reagent, R.T.U. (PK-7100) was applied for 30 min, followed by three PBS washes, ImmPACT DAB substrate (SK-4105) development for 2-3 min, and three additional PBS washes. Sections were coverslipped with VECTASHIELD Antifade Mounting Medium (Vector Laboratories, H-1000-10). Paired fluorescent and brightfield images were acquired on a Keyence BZ-X710 fluorescence microscope.

The following primary antibodies were used: GFP (1:500, Abcam, ab6673), human NCAM (1:500, Santa Cruz, sc-106), human nuclei (rabbit 1:200, NeoBiotechnologies, RBM8-8775-P1ABX; mouse 1:200, Sigma, MAB1281), Iba1 (1:500, Abcam, ab5076), human TMEM119 (1:200, Abcam, ab185333), P2RY12 (1:200, Sigma, HPA014518), NeuN (1:500, Abcam, ab279297), human GFAP (1:500, Takara Bio, STEM123, Y40420), Nestin (1:200, Sigma, MAB5326), SOX1 (1:50, R&D Systems, AF3369), Ki-67 (1:100, Abcam, ab16667), Tau (MC1; 1:1000 Peter Davis, Northwell Health), GFAP (1:500, Abcam, ab4674), and PGRN (1:100, R&D Systems, AF2557).

Secondary antibodies were used at 1:500 for 1 h, including Alexa Fluor 488 Donkey Anti-Goat IgG, Alexa Fluor 568 Donkey Anti-Rat IgG, Alexa Fluor 488 Donkey Anti-Rabbit IgG, Alexa Fluor 568 Donkey Anti-Mouse IgG (Thermo Fisher Scientific), Goat Anti-Chicken IgG Alexa Fluor 488, and Streptavidin Alexa Fluor 568 conjugate (Fisher, S11226).

TUNEL staining was performed after secondary antibody incubation using the In Situ Cell Death Detection Kit, TMR Red (Roche, 12156792910) according to the manufacturer’s instructions by incubating sections in TUNEL reaction mixture (50 µl enzyme solution + 450 µl label solution) in a humidified chamber at 37 °C for 1 h in the dark, followed by three PBS washes and mounting in antifade medium.

### Single-nucleus RNA sequencing (snRNAseq)

Freshly dissected mouse brains were chopped into 400 μm brain slices and put into ice-cold Hanks′ Balanced Salt solution (HBSS; Sigma H6648) using a tissue chopper (McIlwain). Human grafts were micro-dissected under a brightfield dissection microscope based on engraftment location and tissue morphology then snap-frozen on dry ice and stored in −80°C. Nuclei isolation and snRNAseq workflow follows previous studies (*70*). Briefly, micro-dissected human grafts were dissociated in 1.5 ml of Nuclei PURE lysis buffer (Sigma, NUC201-1KT) and homogenized in a Dounce tissue grinder (Sigma, D8938-1SET) using 15 strokes with pestle A followed by 15 strokes with pestle B. The lysate was then passed through a 35 µm strainer (Falcon 352235), then centrifuged at 600g for 5 minutes at 4°C, followed by washing three times with 1 ml PBS containing 1% bovine serum albumin (BSA, Thermo Fisher Scientific, 37525), 20 mM DTT (Thermo Fisher Scientific, 426380500), and 0.2 U/µl recombinant ribonuclease (RNase) inhibitor (Ambion, AM2684). Nuclei were pelleted again at 600g for 5 min at 4°C and resuspended in 350 μl PBS containing 0.04% BSA and DAPI. DAPI-positive nuclei were then sorted by the fluorescence-activated cell sorting (FACS) core at the Weill Cornell Medicine to remove debris, counted, and diluted to 1,000 nuclei/µl in PBS containing 0.04% BSA. cDNA libraries were then generated following Chromium Single Cell 3′ Reagent Kits v3 (10x Genomics, PN-1000075) according to the manufacturer’s instructions. Libraries were sequenced on a NovaSeq 6000 (Illumina) using a 100-cycle run.

### snRNA-seq data processing

Raw sequencing reads were initially aligned to a combined human–mouse reference genome (GRCh38 and GRCm39; refdata-gex-GRCh38_and_GRCm39-2024-A) using cellranger (v9.0.0, 10x Genomics). Species identity for each cell barcode was determined from the gem_classification.csv output. Barcodes classified as human were extracted and retained for downstream analysis. The original dataset was subsequently aligned using a human-only reference genome (refdata-gex-GRCh38-2024-A) with cellranger count and reprocessed using cellranger reanalyze, restricting the analysis to human cell barcodes via the --barcodes option. To better capture nuclear RNA, reads mapping to pre mRNA were included in the count matrix, and default cellranger parameters were used for barcode calling. Genes detected in three or fewer cells were excluded. Nuclei were further filtered to remove those with fewer than 200 or more than 7000 detected genes, more than 30,000 UMIs, or a high fraction of mitochondrial reads (>5%). Potential doublets were identified in each sample separately using DoubletFinder [ref McGinnis 2019 Cell Systms], and high confidence doublets were removed.

Downstream normalization, clustering, and integration were performed in Seurat (v5.0.0). Counts were normalized by total library size (scale factor 10,000) and log transformed. Highly variable genes (n = 2000) were selected using FindVariableFeatures() with the vst method, yielding corrected UMI counts, log transformed expression values, and Pearson residuals from a regularized negative binomial regression model. Principal component analysis was performed, and t distributed stochastic neighbor embedding was computed using the top 15 principal components. Clusters were identified using FindNeighbors (top 15 PCs) and FindClusters (resolution = 0.1). For DoubletFinder, the neighborhood size parameter pK was selected using the mean variance normalized bimodality coefficient (BCmvn) approach, with 15 PCs and pN set to 0.25. Samples were integrated using Seurat’s FindIntegrationAnchors() and IntegrateData() functions. We integrated 4-month post-engrafted HPC+NPC human nuclei from WT(3R) NPC+HPC (n=3), 4R-WT NPC+HPC control (n=4), 4R-WT NPC+HPC+tau seeds (n=4), 4R-P301S NPC+HPC control (n=4), and 4R-P301S NPC+HPC tau seeds (n=5).

### RNA velocity based pseudotime

To get the RNA velocity based pseudotime per cell, we followed the tutorial of Velocyto 0.17 (*71*) and scVelo 0.3.1 (*72*). Loom files were generated by running Velocyto 0.17 for all 3R and 4R wild-type samples (3 and 4 samples respectively). After reading in the spliced and unspliced counts stored in loom files, preprocessing was performed by setting min_shared_counts=10 and n_top_genes=3000. Moments for velocity estimation were obtained by setting n_pcs=30 and n_neighbors=30, which was followed by RNA velocity estimation. Velocity pseudotime and latent time were computed based on the velocity graph.

### DEG and pathway analysis

Differential gene expression analysis for each cell type was performed using the FindMarkers function in Seurat with the MAST method (*73*) and the parameter “min.cells.feature = 10”. Genes with an absolute log2 fold change >0.1 and an adjusted p value <0.05 were considered differentially expressed genes. This relatively permissive log2 fold change threshold was used to preserve sensitivity for detecting transcriptional changes in vulnerable or rare neuronal populations. Because tau seeded 4R P301S grafts undergo neuronal loss, highly stringent DEG filtering could preferentially exclude small populations of neurons transitioning toward degenerative or cell death associated states. Normalized nuclei counts were generated using the dittoSeq R package (*74*). Gene Ontology and pathway enrichment analyses were conducted using the pathfindR and clusterProfiler packages in R as previously described (*75, 76*). ChatGPT 5.5 was used to assist with generating R analysis code and organizing the structure of R scripts. DEGs were also analyzed using Ingenuity Pathway Analysis (IPA; Qiagen). For IPA, DEG tables were uploaded with average log2 fold change assigned as the expression log ratio and adjusted p value assigned as the experimental p value. IPA core expression analysis was performed using the input expression log ratio to infer enriched canonical pathways, upstream regulators, and disease or functional annotations.

Microglial maturity gene list (*27*): *SPP1, CD74, ACTB, C3, FTL, FOS, CSF1R, B2M, C1QC, C1QB, PSAP, A2M, ITM2B, LAPTM5, CTSB, P2RY12, C1QA, SLCO2B1, RGS1, APOE, CCL4L2, RNASET2, NEAT1, CX3CR1, DUSP1, SAT1, ZFP36, CD81, HLA-B and HLA-DRA*.

Astrocyte maturity gene list (*31*): *SLC1A2, SPARCL1, SLC1A3, GLUL, CPE, GJA1, ATP1B2, AQP4, GPM6A, ATP1A2, and GLUD1*.

### Golgi-Cox staining and imaging

Because Golgi-Cox impregnation requires fresh, unfixed tissue, staining was performed on a separate cohort of mice (n=4) transplanted with 4R-WT NPCs and HPCs using the FD Rapid GolgiStain Kit (FD NeuroTechnologies, PK401A), as previously described with minor modifications (*74*). Briefly, equal volumes of Solutions A and B were combined at least 24 h before perfusion. Mice were deeply anesthetized and transcardially perfused with phosphate-buffered saline (PBS). Brains were rapidly dissected and processed per the manufacturer’s instructions: impregnation in the combined A/B solution for two weeks, followed by incubation in Solution C for three days. Brains were then snap-frozen in dry-ice–cooled isopentane and sectioned at 100 µm on a cryostat (Leica). Sections containing the human graft were mounted on gelatin-coated slides (FD NeuroTechnologies, PO101), air-dried overnight, developed in diluted Solutions D and E, dehydrated, and coverslipped in Permount mounting medium (Fisher, SP15-500) according to the kit manual.

Golgi-stained sections were imaged under brightfield illumination on a Keyence BZ-X710 all-in-one fluorescence microscope. For whole-neuron reconstructions, z-stacks were acquired with a 40X or 63X objective at 0.7 µm step size and stitched to capture the full neuronal arbor. For dendritic spine and branch imaging, z-stacks spanning the full depth of individual dendrites were acquired with a 100× objective at 0.1 µm step size using the multi-stack module. Stitched image sets were processed in ImageJ by thresholding, followed by manual tracing and quantification of neurites as previously described (*77*).

### Spatial transcriptomics

Immunostaining in combination with Visium HD spatial transcriptomics was performed using the 10x Visium HD kit (10x Genomics, 1000675) according to the manufacturer’s instructions, with minor modifications. Mouse brains were dissected as described above, embedded in O.C.T. compound (Sakura, Tissue Tek O.C.T. 4583), snap frozen on dry ice, sealed, and stored at −80°C. Embedded brains were cryosectioned at 10 µm thickness and sections were stored at −80°C until use. Individual sections were mounted in the center of Visium HD glass slides. Immunofluorescence staining was performed as previously described (*78*). Briefly, slides were removed from −80°C and dried at 37°C for 1 min on a 10x Genomics thermocycler adapter placed in a thermocycler (Bio-Rad). Sections were fixed in precooled 100% methanol at −20°C for 5 min, washed twice with 3x saline sodium citrate buffer (3x SSC; Sigma, S6639 1L), and loaded into a slide cassette. Sections were blocked for 20 min in 10% BSA (Miltenyi Biotec, 130091376) supplemented with RNase inhibitor (1 U/µL; Ambion, AM2684). Primary antibodies were diluted in the same blocking solution and applied to sections for 30 min at room temperature. Tau (MC1; Peter Davis; Northwell Health) 1:1000; human nuclei antibody 1:200 (NeoBiotechnologies, RBM8-8775-P1ABX). Slides were washed three times with 3x SSC, followed by incubation with secondary antibodies and DAPI diluted in 10% BSA plus RNase inhibitor for 15 min. Slides were then washed four times and rinsed extensively with 3x SSC. Sections were mounted in an RNase compatible mounting medium containing 85% glycerol, 10% RNase inhibitor, and 5% water, immediately imaged at 20x on a Keyence microscope with tile stitching, and then submitted to the Epigenomics Core at Weill Cornell Medicine for Visium CytAssist HD processing, library preparation, and sequencing following the 10x Genomics manual.

Visium HD spatial transcriptomic libraries were prepared and sequenced according to the Visium HD Spatial Gene Expression Reagent Kits User Guide (CG000685, 10x Genomics). Raw sequencing data were processed using Space Ranger v4.0.1 through the 10x Genomics Cloud Analysis platform with default parameters. Reads were aligned to the GRCh38 human reference genome to generate spatially resolved gene expression matrices. The 8 µm binned output was used for all downstream spatial analyses. Loupe Browser v9.0.0 was used to visualize the 8 µm binned data, manually annotate regions of interest, and export corresponding barcodes. Gene lists of interest were imported into Loupe Browser through the Features tab. IFN high spatial regions were defined as barcodes with log2 feature sum expression greater than 2 for the IFN upregulated gene set, and exported for overlay with immunostaining images and cell death gene heatmaps. To assess the spatial distribution of seed induced HNRNPK-AS1 associated transcription factors across cell death enriched domains, barcodes were stratified into cell death high and cell death low regions based on log2 feature sum expression greater than or less than 3.2 for the cell death gene set. Expression distributions of seed induced HNRNPK-AS1 associated transcription factors were then analyzed in Loupe Browser and visualized as violin plots.

### Electrophysiology

Following isoflurane anesthesia, mice were decapitated, and brains were rapidly extracted into ice-cold oxygenated cutting solution containing 95% O₂ and 5% CO₂. The cutting solution contained, in mM: 212.5 sucrose, 10 D-glucose, 3 KCl, 1.25 Na₂HPO₄, 24 NaHCO₃, 0.1 CaCl₂, and 5 MgCl₂, pH 7.4. Horizontal brain slices containing the hippocampus were prepared at 300 µm thickness using a vibratome (Campden) in the same cutting solution. Slices were then transferred to oxygenated artificial cerebrospinal fluid (ACSF) containing, in mM: 130 NaCl, 3.5 KCl, 1.25 NaH₂PO₄, 24 NaHCO₃, 10 D-glucose, 1.5 CaCl₂, and 1.5 MgCl₂, pH 7.4. Slices were incubated for at least 30 min at 32°C, followed by 1 h at room temperature before recording. Individual slices were transferred to an immersion recording chamber and continuously submerged in oxygenated ACSF. Whole-cell patch-clamp recordings were performed at room temperature, 20 to 22°C. Neurons were identified in DIC mode using a ZEISS Axio Examiner upright microscope equipped with an ORCA Fusion Digital CMOS high-resolution camera (Hamamatsu). Whole-cell membrane currents were amplified using a Multiclamp 700B amplifier, and data acquisition was controlled by pCLAMP 11.3 software (Molecular Devices). A Digidata 1550B interface was used for digital-to-analog and analog-to-digital signal conversion between the amplifier and computer. Data were sampled at 10 kHz and filtered at 2 kHz. Recording pipettes, 6 to 8 MΩ, were fabricated from borosilicate glass capillaries (1B150F-4, World Precision Instruments) using a PC-100 vertical pipette puller (Narishige). Pipettes were filled with internal solution containing, in mM: 130 K-gluconate, 10 HEPES, 5 NaCl, and 0.2 EGTA, supplemented with 0.2% biocytin, with pH adjusted to 7.3 using KOH. Action potentials were recorded in current-clamp mode by applying 500 ms step current injections in 20 mV increments. Putative excitatory and inhibitory neurons were distinguished based on soma size and established differences in intrinsic firing properties, including spike frequency adaptation, firing rate, and action potential waveform, as previously described.(*79–83*) Spontaneous excitatory postsynaptic currents were recorded in voltage-clamp mode at a holding potential of −60 mV. Resting membrane potential was measured in I = 0 mode. Membrane properties, including cell membrane capacitance (Cm), membrane resistance (Rm), and access resistance (Ra), were monitored using the Membrane Test function in pCLAMP. Only neurons with Ra <40 MΩ at break-in and with Ra remaining within 20% of the initial value throughout the recording were included in the analysis. Offline analysis was performed using Clampfit version 11.3 and NeuroExpress version 25.4.23.

### Single-nuclei isoform RNA sequencing (SnISOr-Seq)

SnISOr Seq libraries were generated as previously described (*84*), with minor modifications. Briefly, 10x Genomics Chromium Single Cell 3′ barcoded cDNA was first subjected to a linear, asymmetric PCR enrichment using the Partial Read1 primer to preferentially amplify barcode containing molecules, followed by 0.8× SPRIselect cleanup. A second exponential PCR was then performed using Partial Read1 and Partial TSO primers, followed by 0.6× SPRIselect cleanup and elution in EB buffer. KAPA HiFi HotStart ReadyMix was used for all amplification steps unless otherwise stated. Library size and quality were assessed using Agilent TapeStation Genomic DNA ScreenTape.

To enrich for spliced cDNA molecules, an exome capture step was performed using SureSelectXT HSQ reagents with the SureSelectXT HS Human All Exon V8 probe set (Agilent, 5191-6873), following the manufacturer’s instructions. Captured molecules were recovered on M-270 streptavidin Dynabeads, washed under stringent conditions, and PCR amplified with Partial Read1 and Partial TSO primers before a final 0.6× SPRIselect cleanup. For long read sequencing, approximately 75 fmol of enriched cDNA per sample was used for Oxford Nanopore library preparation with the Ligation Sequencing Kit (SQK LSK110). Libraries were sequenced on PromethION flow cells (FLO PRO002) for up to 72 h. Basecalling was performed using Guppy with a minimum Q score threshold of 9, and reads were aligned to the human reference genome (GRCh38).

PCR induced molecular duplication was corrected by collapsing reads using UMIs. For UMI groups mapping to the same locus, reads containing UMIs within an edit distance of 4 from a more abundant UMI were discarded. Alternative exon inclusion and exclusion events were quantified for each cell type and experimental condition using scisorATAC (*72*). Inclusion and exclusion counts were summarized as 2 × 2 contingency tables for differential exon usage testing by Fisher’s exact test. Multiple hypothesis testing was controlled using the Benjamini and Yekutieli procedure. Exons were retained for downstream analyses if Ψ values fell between 5% and 95% and were supported by at least one read per condition. SnISOr Seq outputs were parsed using custom Python scripts to extract Ψ, ΔΨ, and associated probabilities of splicing change for each exon in each cell type. Events were prioritized for visualization if |ΔΨ| exceeded 0.1. No-seed WT-HPC, n = 3 mice; seeded WT-HPC, n = 3 mice; PGRN-iso, n = 3 mice.

### HNRNPK-AS1 transcription factor (TF) motif analysis

Predicted transcription factor binding motifs within the HNRNPK-AS1 promoter were identified using FIMO from the MEME Suite. The HNRNPK-AS1 promoter region was defined as GRCh38/hg38 chr9:83,956,132-83,957,209 on the positive strand, corresponding to the proximal HNRNPK-AS1 promoter and transcription start region based on UCSC Genome Browser annotation. The promoter sequence was formatted as a FASTA file in R and scanned against the JASPAR2026 CORE vertebrate nonredundant position frequency matrix database in MEME format. FIMO was run using a p value threshold of 1 × 10^-4^ with both DNA strands scanned. The resulting motif level output was imported into R, where predicted motif hits were mapped to transcription factor gene names and cross compared with the PGRN OE rescued DEG list. This analysis identified candidate HNRNPK-AS1 regulatory transcription factors whose expression was altered by tau seeding and restored by microglial PGRN OE. The rescued transcription factor list was then used for NicheNet analysis to infer candidate microglia to neuron signaling relationships (*48*). The NicheNet 2.0 human ligand target matrix and prior networks were used as reference databases (*49*). Predicted ligand to transcription factor links were filtered using a regulatory potential threshold corresponding to the top 5% of NicheNet weights (weight_quantile_cutoff = 0.95). For visualization, the top five transcription factor links per ligand were retained (top_links_per_ligand = 5) and plotted to summarize the strongest predicted microglial ligand to neuronal transcription factor signaling relationships associated with the HNRNPK-AS1 regulatory program.

### Lentivirus generation and transduction

HNRNPK-shRNA (Sigma, TRCN0000295992, TGATCTTGGTGGACCTATTAT) and control shRNA lentivirus (Sigma, SHC016V) were purchased from Sigma. Lentiviral particles for HNRNPK-AS1 gRNAs and none-target control (NT) sgRNA were generated as previously described (*14*). Briefly, HEK293T cells (ATCC; Cat. No. CRL-11268) were maintained in DMEM/F-12 supplemented with 10% FBS and 1% penicillin/streptomycin and seeded into 10 cm dishes to reach the recommended confluency at transfection. Lentiviral vectors were produced by co transfecting transfer plasmid together with third generation packaging plasmids pMDLg/pRRE (Addgene plasmid #12251), pRSV-REV (Addgene plasmid #12253), and pMD2.G (Addgene plasmid #12259) using Lipofectamine™ 3000 Reagent (Thermo Fisher Scientific; Cat. No. L3000015) in Opti-MEM I Reduced Serum Medium (Thermo Fisher Scientific; Cat. No. 31985070), following the manufacturer’s protocol. Viral supernatant was harvested at defined time points, pooled, clarified, and filtered through a 0.45 μm PVDF filter, then concentrated using Lenti-X Concentrator (Takara Bio; Cat. No. 631232), and resuspended in sterile DPBS (Thermo Fisher Scientific; Cat. No. 14190144) and aliquoted for storage at −80°C. For lentiviral transduction, differentiated 4R-P301S neurons were exposed to lentivirus at an estimated MOI of 3 in culture medium. The medium was replaced the following day.

### Cell viability test

4R-P301S neurons were cultured for 2 weeks following lentiviral HNRNPK-shRNA transduction. Culture medium was replaced with PBS containing 1ug/ml DAPI (Fisher, EN62248) and 1ug/ml Calcein AM (Invitrogen, C3099), and cells were incubated for 5 min before imaging using a Keyence microscope. Cells were then carefully washed three times with PBS and processed for the CellTiter-Glo Cell Viability Assay (Promega, G9241) according to the manufacturer’s instructions. Three independent differentiations with lentivirus-mediated HNRNPK knockdown were included in this study.

### Bulk RNA-sequencing and data analysis

Three independent differentiation and lentivirus mediated HNRNPK knockdown was included in this study. For each differentiation, cells were lysed and RNA was isolated using RNeasy Plus Mini Kit (Qiagen, 74134) following manufacture’s instruction. RNA concentration was determined using Nanodrop Spectrophotometer (Thermo scientific). Total RNA was sequenced using Novogene EukmRNA Seq, Directional with PolyA service with data analysis. Differentially expressed gene (DEG) analysis was further conducted using the DESeq2 package in R(*85*). Genes that have absolute log2 fold change >0.1 and adjusted p value <0.05 are considered DEGs. Modeling Alternative Junction Inclusion Quantification (MAJIQ V3) splicing analysis pipeline was conducted as previously described (*86*). Briefly, MAJIQ was performed in a Linux environment using Windows Subsystem for Linux (WSL). Paired end FASTQ files were aligned to the human reference genome (GRCh38 primary assembly) with STAR, using a genome index built with GENCODE v45 annotations and generating coordinate sorted BAM files. MAJIQ v3 was then used to construct splice graphs with the MAJIQ build pipeline using the GENCODE v45 annotation (GFF3) and an experiment design table listing BAM paths and sample groups (3 Ctrl and 3 KD replicates), producing a splicegraph.zarr and junction files for each sample. Per sample splice junction usage was quantified with majiq psi coverage (minreads 10, minbins 3) to generate psi files. Differential splicing between KD and Ctrl was assessed using MAJIQ heterogen and MAJIQ deltapsi with replicate aware comparisons, and outputs were generated both as MAJIQ summary tables and VOILA compatible files. For visualization and reporting, group level splice graph coverage files were created with sg coverage and VOILA was used to export annotated TSV tables and to review events interactively.

### RT-qPCR

Leftover RNAs from bulk RNAseq were also processed for RT-qPCR to confirm the knockdown of HNRPNK. cDNA was generated using the iScript Reverse Transcription Supermix for RT-qPCR (Bio-Rad; Cat. No. 1708840), and qPCR was performed using SsoAdvanced Universal SYBR Green Supermix (Bio-Rad; Cat. No. 1725270) with primers listed below. Thermal cycling was carried out at 95°C for 10 min, followed by 45 cycles of 95°C for 15 s, 60°C for 30 s, and 72°C for 30 s. Data were acquired on the BCFX384 Touch Real-Time PCR Detection System (Bio-Rad). Each sample was run in replicate, and relative gene expression was calculated using the ΔΔCT method.

*18s*_for 5’-AGTCCCTGCCCTTTGTACACA-3’,
*18s*_rev 5’-GATCCGAGGGCCTCACTAAAC-3’.
HNRNPK forward: 5’-CGCCCTGCAGAAGATATGGA-3’,
HNRNPK reverse: 5’-CCTTTTCCAATCACTGCCCC-3’

### RNA scope

RNAscope in situ hybridization was performed using the ACD RNAscope Multiplex Fluorescent Reagent Kit v2 (Advanced Cell Diagnostics, Bio-Techne) following the manufacturer’s protease-free protocol optimized for fixed-frozen tissue. Briefly, cryosections were post-fixed in 4% paraformaldehyde, dehydrated through graded ethanol, and air-dried before probe hybridization. Target probes (7ZZ probe, Hs-HNRNPK-AS1-No-XMm) were hybridized for 2 h at 40 °C in the HybEZ oven, followed by sequential amplification steps using AMP reagents and detection with Opal fluorophores. Slides were counterstained with DAPI and mounted in antifade medium.

### Statistics

Statistical analyses were performed using GraphPad Prism 11. Unless otherwise indicated in the figure legends, comparisons between two groups were performed using unpaired two-tailed t tests, and comparisons among more than two groups were performed using one-way ANOVA followed by Tukey’s multiple-comparisons test. For fig. S5F, a two-tailed Mann-Whitney U test was used because one group showed high variability. For fig. S13B and S13D, two-tailed Welch’s t tests were used to account for unequal variances between groups. For Fig. 6C and Fig. 6E, one-way ANOVA followed by Dunnett’s multiple-comparisons test was used to compare each condition with the tau-seeded WT-HPC control group. Exact statistical tests are indicated in the corresponding figure legends.

## Acknowledgments

We thank the Epigenetics Core for Visium HD library preparation and sequencing. Illustrations were created with BioRender.com.

## Author contributions

Conceptualization: WQ, LG

Methodology: WQ, LF, MWJ, PY, EC, AA, RKN, SG, LG

Investigation: WQ, LF, MWJ, PY, EC, AA, MYW, WL, SG, LG

Data curation: WQ, LF, AA, WH

Formal analysis: WQ, LF, MWJ, EC, AA, WH

Validation: WQ, EC, SG

Resources: RKN, MBJ, SG

Supervision: HUT, AO, SG, LG

Funding acquisition: WQ, LG

Project administration: WQ, LG

Visualization: WQ, LF, MWJ, AA, WH, SG

Writing-original draft: WQ, LG

Writing-review & editing: WQ, LF, MWJ, PY, EC, AA, WH, RKN, MYW, WL, MBJ, HT, AO, SG, LG

## Data, code, and materials availability

Bulk (GSE335751), single-cell RNA-seq data (GSE335853), and spatial transcriptomic data (GSE335784) have been deposited at GEO and are publicly available as of the date of publication. Any additional information required to reanalyze the data reported in this paper is available from the lead contact upon reasonable request.

---

**Fig. S1.**
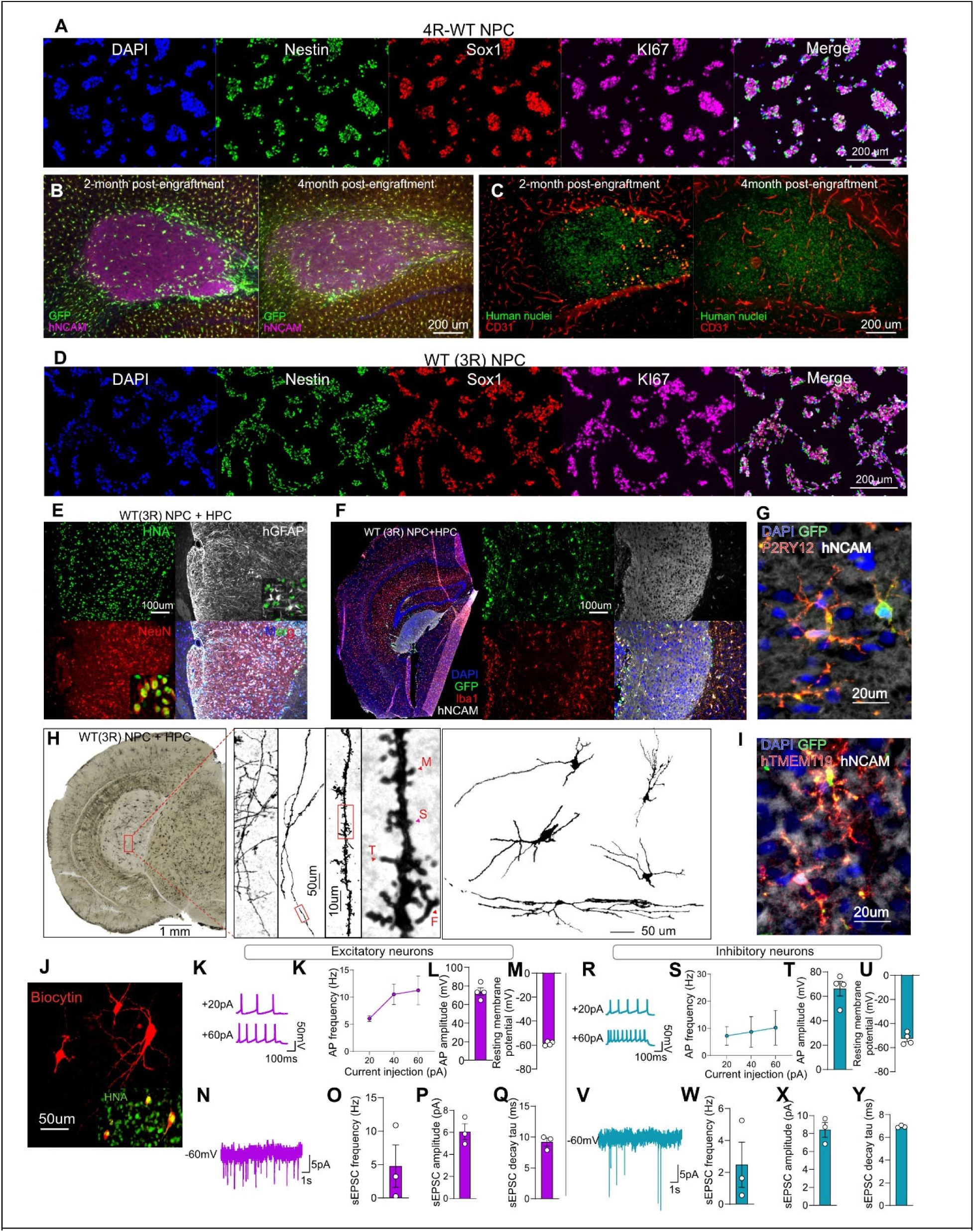
Additional characterization of HuMiNAX. (**A**) Immunostaining of 4R-WT NPCs showing expression of the neural progenitor markers Nestin and SOX1, together with the proliferation marker KI67. (**B**) Representative images at 2 and 4 months post engraftment showing GFP⁺ human microglia distributed within hNCAM⁺ human neural grafts. (**C**) Representative images at 2 and 4 months post engraftment showing vascularization of human nuclei⁺ graft tissue, visualized by CD31 staining. (**D**) Immunostaining of WT(3R) NPCs showing expression of Nestin, SOX1, and KI67. (**E**) Immunostaining of WT(3R) NPC + HPC grafts showing differentiation of human graft cells into HNA⁺ NeuN⁺ neurons and hGFAP⁺ astrocytes. (**F**) Immunostaining showing that co transplanted HPCs generate GFP⁺ IBA1⁺ human microglia that populate hNCAM⁺ human neural grafts. (**G**) High magnification image showing graft associated human microglia expressing the homeostatic microglial marker P2RY12. (**H**) Golgi Cox staining of grafted WT(3R) human neurons four months after transplantation, showing dendritic arborization, dendritic spines of multiple morphologies, and representative neuronal reconstructions. (**I**) High magnification image showing graft associated human microglia expressing the homeostatic microglial marker TMEM119. (**J**) Immunostaining of a biocytin filled recorded neuron confirming human origin by HNA labeling. (**K-M**) Representative action potential traces and intrinsic membrane properties of graft derived excitatory neurons, including action potential frequency, action potential amplitude, and resting membrane potential. N = 4 mice. (**N-Q**) Representative spontaneous excitatory postsynaptic current recording from graft derived excitatory neurons, with quantification of sEPSC frequency, amplitude, and decay tau. N = 3 mice. (**R-U**) Representative action potential traces and intrinsic membrane properties of graft derived inhibitory neurons, including action potential frequency, action potential amplitude, and resting membrane potential. N = 4 mice. (**V-Y**) Representative spontaneous excitatory postsynaptic current recording from graft derived inhibitory neurons, with quantification of sEPSC frequency, amplitude, and decay tau. N = 3 mice. Data are shown as mean ± SEM. For electrophysiological quantifications, each data point represents the average value from one mouse.

**Fig. S2.**
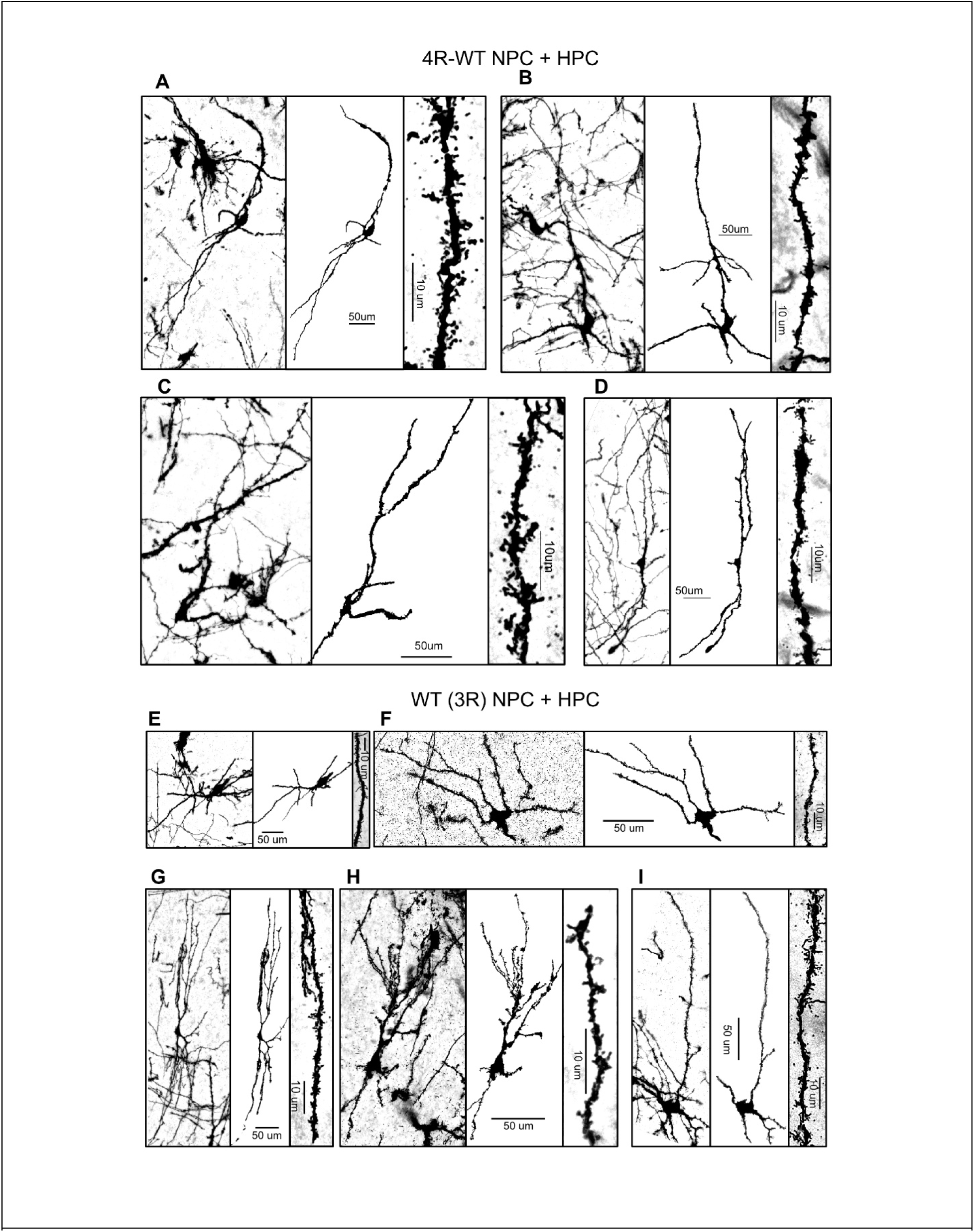
Golgi-Cox staining, neuronal reconstructions, and dendritic spine morphology in HuMiNAX grafts. (**A-D**) Representative Golgi-Cox-stained neurons, corresponding neuronal reconstructions, and high-magnification dendritic branch images from grafts generated by 4R-WT NPC + HPC transplantation, showing complex dendritic arborization and dendritic spine formation. (**E-I**) Representative Golgi-Cox-stained neurons, corresponding neuronal reconstructions, and high-magnification dendritic branch images from grafts generated by WT (3R) NPC + HPC transplantation, showing comparable dendritic maturation and spine morphology. Scale bars are as indicated.

**Fig. S3.**
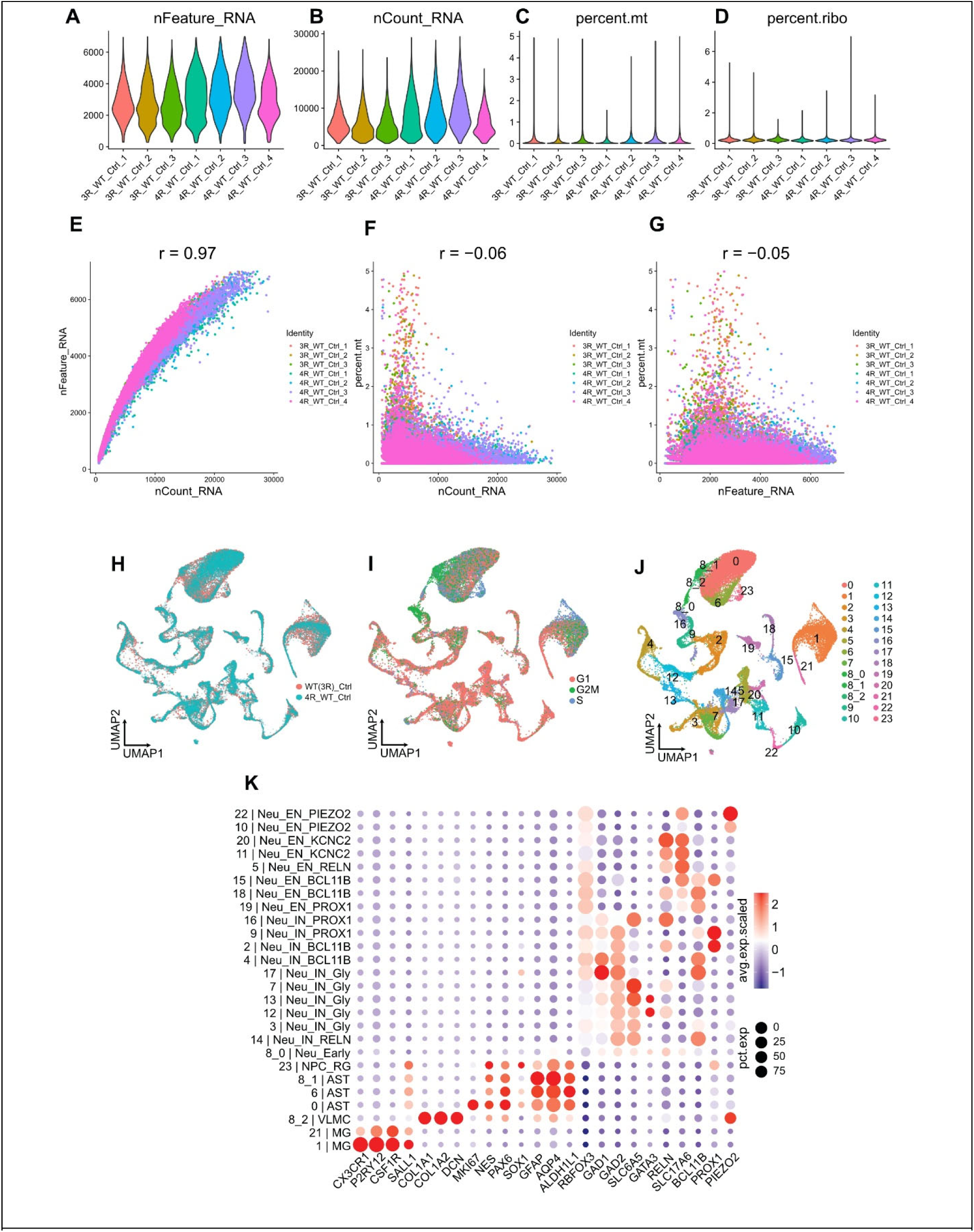
snRNA-seq quality control and cell-type annotation of HuMiNAX datasets. (**A-D**) Violin plots showing per-sample distributions of key snRNA-seq quality control metrics, including the number of detected genes per nucleus, nFeature_RNA (A), total UMI counts per nucleus, nCount_RNA (B), mitochondrial transcript fraction, percent.mt (C), and ribosomal transcript fraction, percent.ribo (D). (**E-G**) Scatter plots showing relationships among quality control metrics across individual nuclei, including nCount_RNA versus nFeature_RNA (E), nCount_RNA versus percent.mt (F), and nFeature_RNA versus percent.mt (G). Each dot represents one nucleus and is colored by sample identity. Pearson correlation coefficients are indicated above each plot. (**H**) UMAP embedding of integrated snRNA-seq datasets showing broad overlap between WT (3R) NPC + HPC and 4R-WT NPC + HPC graft conditions across major cell-type clusters. (**I**) UMAP visualization of inferred cell-cycle phase assignments across all nuclei. (**J**) UMAP showing cluster refinement after subclustering, with cluster 8 subclustered at resolution 0.6 and all other clusters subclustered at resolution 0.3. (**K**) Dot plot summarizing expression of canonical marker genes used for cluster annotation. Cluster numbers and assigned cell-type identities are shown on the y axis, with dot size representing the percentage of nuclei expressing each gene and color indicating scaled average expression.

**Fig. S4.**
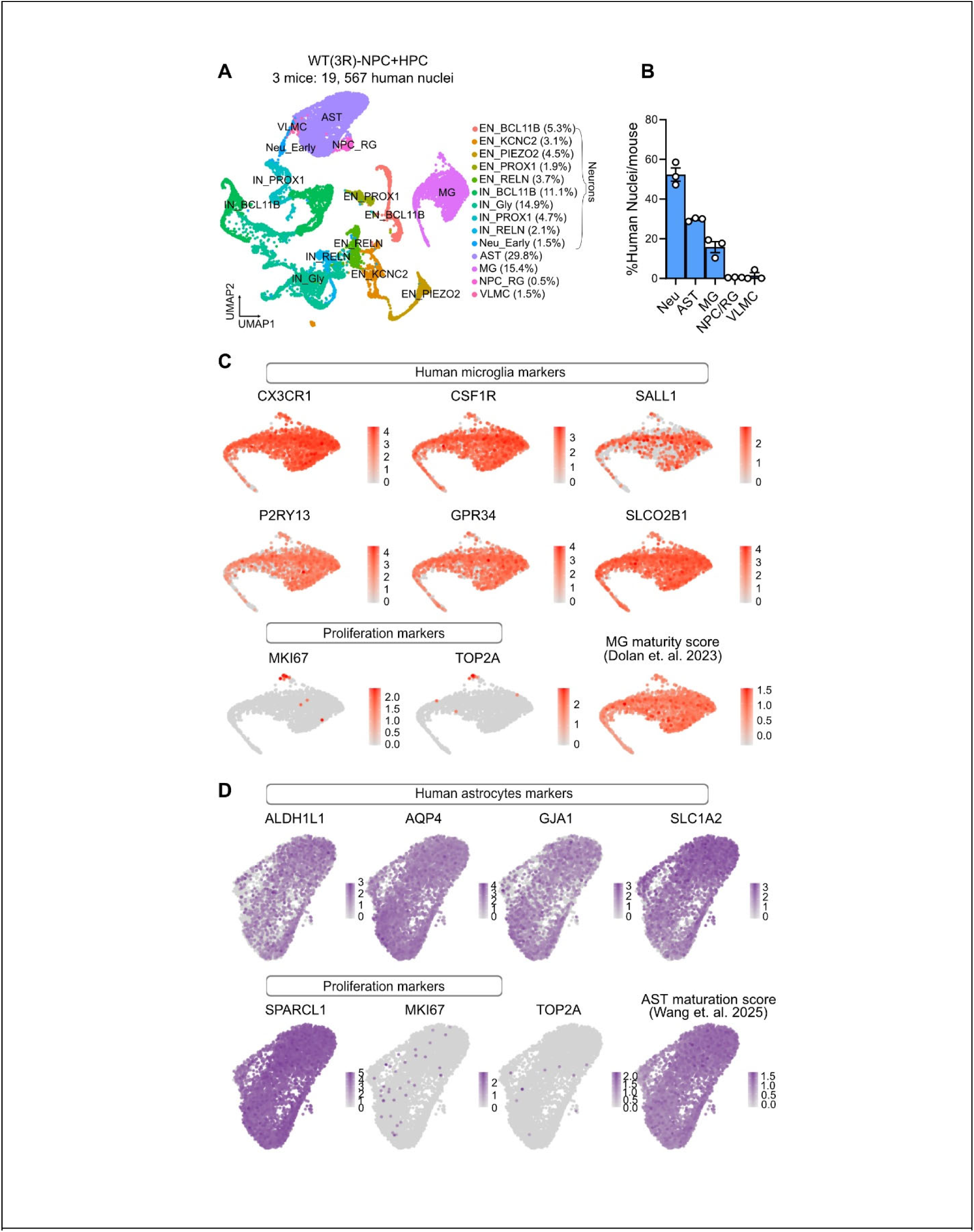
Single-nucleus profiling of glial maturation in WT (3R) HuMiNAX grafts. **(A)** UMAP visualization of snRNA-seq from micro-dissected WT (3R) NPC + HPC grafts, showing 19,567 human nuclei recovered from three mice and annotated into neuronal, astrocytic, microglial, NPC/radial glia, and VLMC populations. (**B**) Quantification of the proportional representation of major human cell classes per mouse, including neurons, astrocytes, microglia, NPC/radial glia, and VLMCs. Data are shown as mean ± SEM. Each dot represents one mouse. N=3 mice. (**C**) Feature plots showing expression of canonical human microglial markers CX3CR1, CSF1R, SALL1, P2RY13, GPR34, and SLCO2B1 in graft-derived microglia, low expression of proliferation markers MKI67 and TOP2A, and enrichment of a human microglial maturity score (*27*). (**D**) Feature plots showing expression of human astrocyte identity markers ALDH1L1, AQP4, GJA1, SLC1A2, and SPARCL1 in graft-derived astrocytes, low expression of proliferation markers MKI67 and TOP2A, and enrichment of an astrocyte maturation score (*31*).

**Fig. S5.**
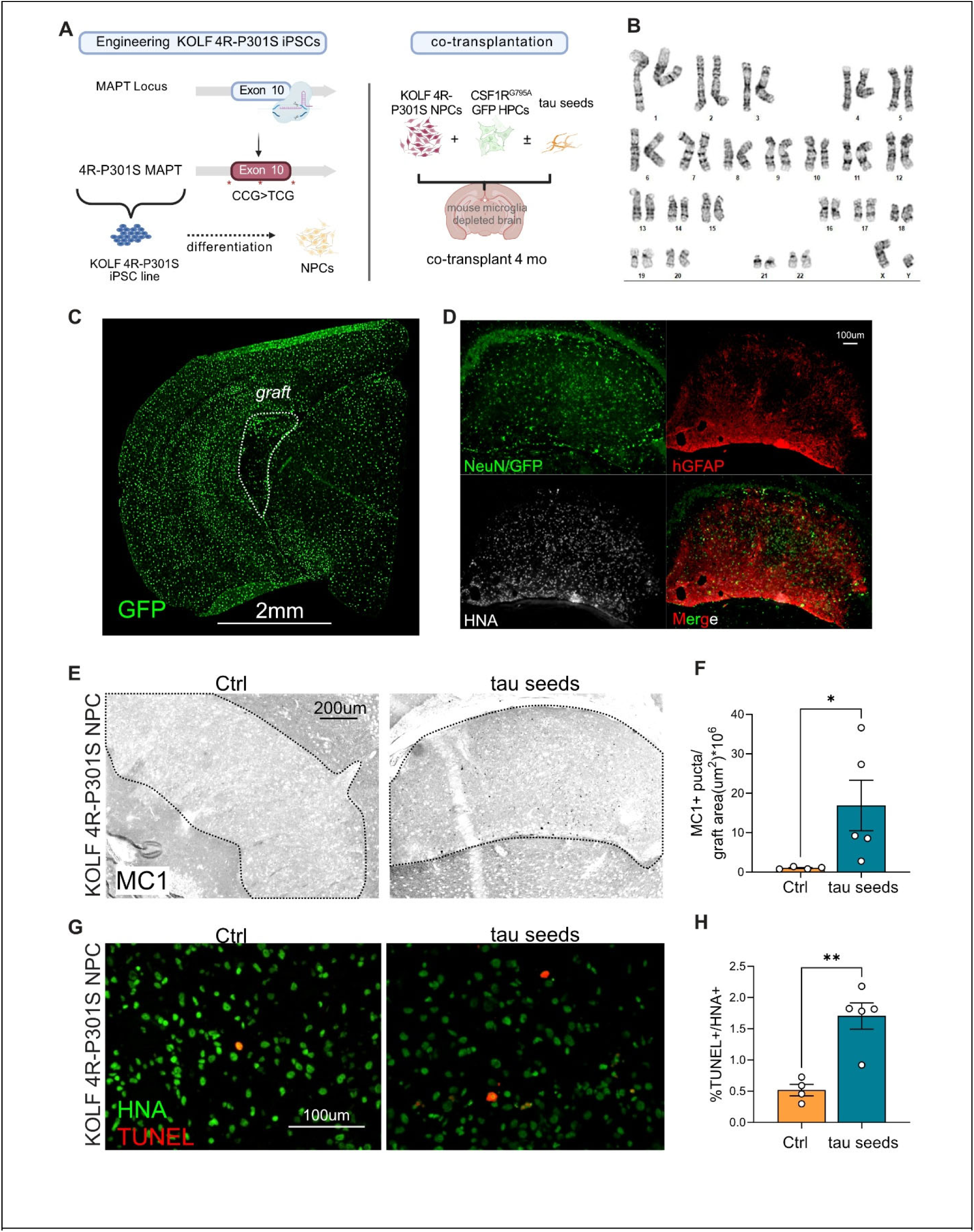
Tau seeding induces pathological tau accumulation and human graft cell death in an independent KOLF 4R P301S iPSC-derived HuMiNAX model. (**A**) Schematic of the generation and transplantation strategy for an independent KOLF 4R P301S iPSC line. CRISPR editing was used to introduce the 4R-P301S into exon 10 of the endogenous MAPT locus. KOLF 4R P301S neural progenitor cells were co-transplanted with CSF1R^G795A^ GFP-labeled human HPC into mouse microglia-depleted adult brains, with or without tau seeds, and analyzed 4 months after transplantation. (**B**) Karyotyping result of the engineered KOLF 4R P301S iPSC line. (**C**) Low-magnification GFP image showing GFP+ human microglia populate the host brain. Dashed outline indicates the graft region. (**D**) Representative immunofluorescence images showing human graft-derived neurons and microglia labeled by NeuN/GFP, human astrocytes labeled by hGFAP, and human graft nuclei labeled by HNA. (**E**) Representative MC1 immunostaining in unseeded control and tau-seeded KOLF 4R P301S grafts. Tau seeding induced MC1-positive pathological tau accumulation within the human graft. Dashed outlines indicate graft regions. (**F**) Quantification of MC1-positive puncta normalized to graft area, showing increased pathological tau burden in tau-seeded KOLF 4R P301S grafts. N=4-5 mice/group; two-tailed Mann-Whitney test. *P < 0.05 (**G**) Representative immunofluorescence images of HNA and TUNEL staining in unseeded control and tau-seeded KOLF 4R P301S grafts. (**H**) Quantification of TUNEL-positive human graft cells, measured as the percentage of TUNEL-positive cells among HNA-positive nuclei. Tau seeding increased human graft cell death in KOLF 4R P301S grafts. two-tailed unpaired t test. **P < 0.01. Data are presented as mean ± SEM, with each dot representing one mouse.

**Fig. S6.**
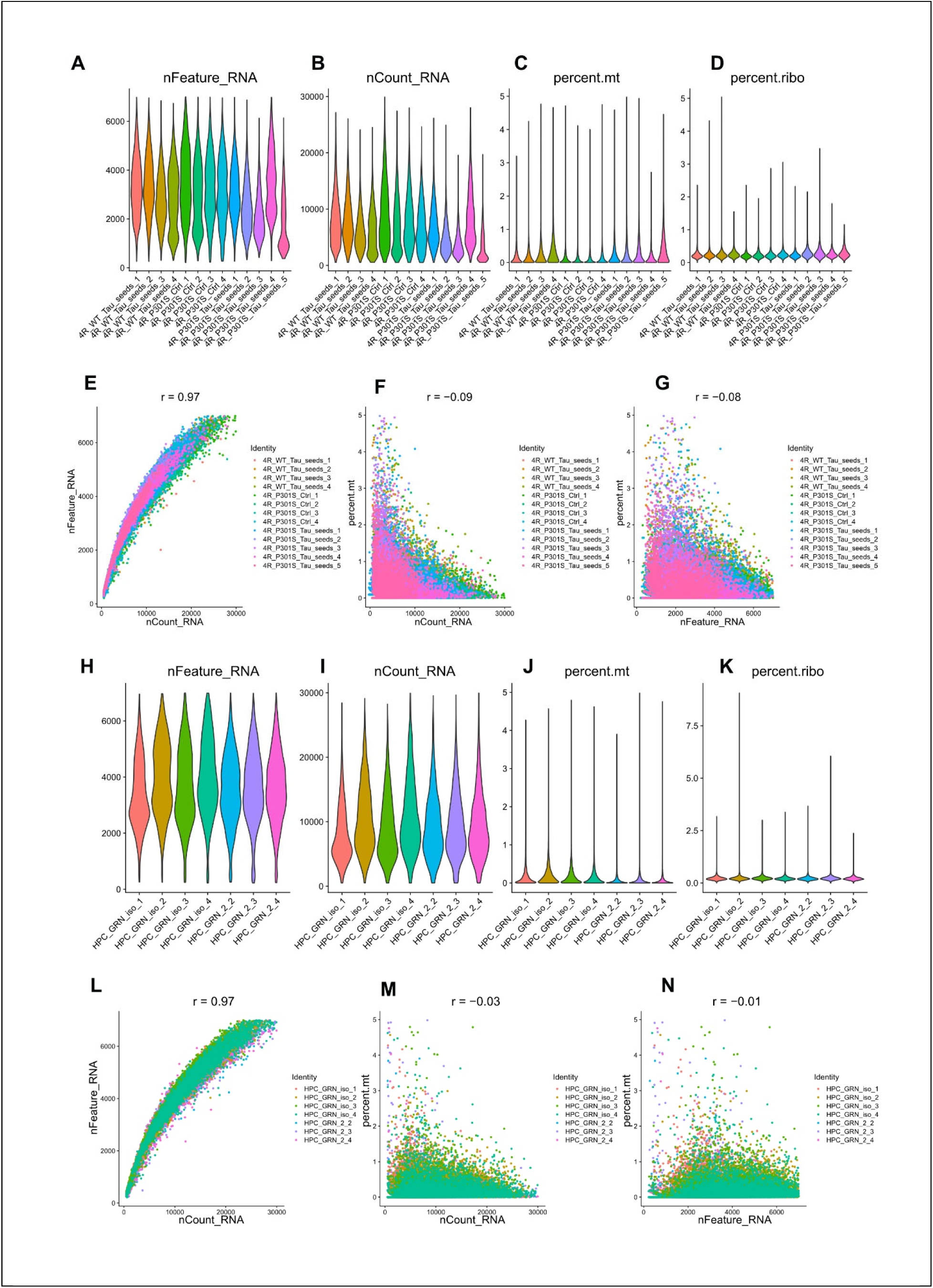
snRNA-seq quality control for 4R tau-seeding and GRN overexpression datasets. (**A-D**) Violin plots showing per-sample distributions of key quality control metrics in the 4R tau-seeding dataset related to Fig. 3, including the number of detected genes per nucleus, nFeature_RNA (A), total UMI counts per nucleus, nCount_RNA (B), mitochondrial transcript fraction, percent.mt (C), and ribosomal transcript fraction, percent.ribo (D). (**E-G**) Scatter plots showing relationships among quality control metrics across individual nuclei in the 4R tau-seeding dataset: nCount_RNA versus nFeature_RNA (E), nCount_RNA versus percent.mt (F), and nFeature_RNA versus percent.mt (G). (**H-K**) Violin plots showing the same quality control metrics in the GRN overexpression dataset related to Fig. 5, including nFeature_RNA (H), nCount_RNA (I), percent.mt (J), and percent.ribo (K). (**L-N**) Scatter plots showing relationships among quality control metrics across individual nuclei in the GRN overexpression dataset: nCount_RNA versus nFeature_RNA (L), nCount_RNA versus percent.mt (M), and nFeature_RNA versus percent.mt (N). Each dot represents one nucleus and is colored by sample identity. Pearson correlation coefficients are shown above each scatter plot.

**Fig. S7.**
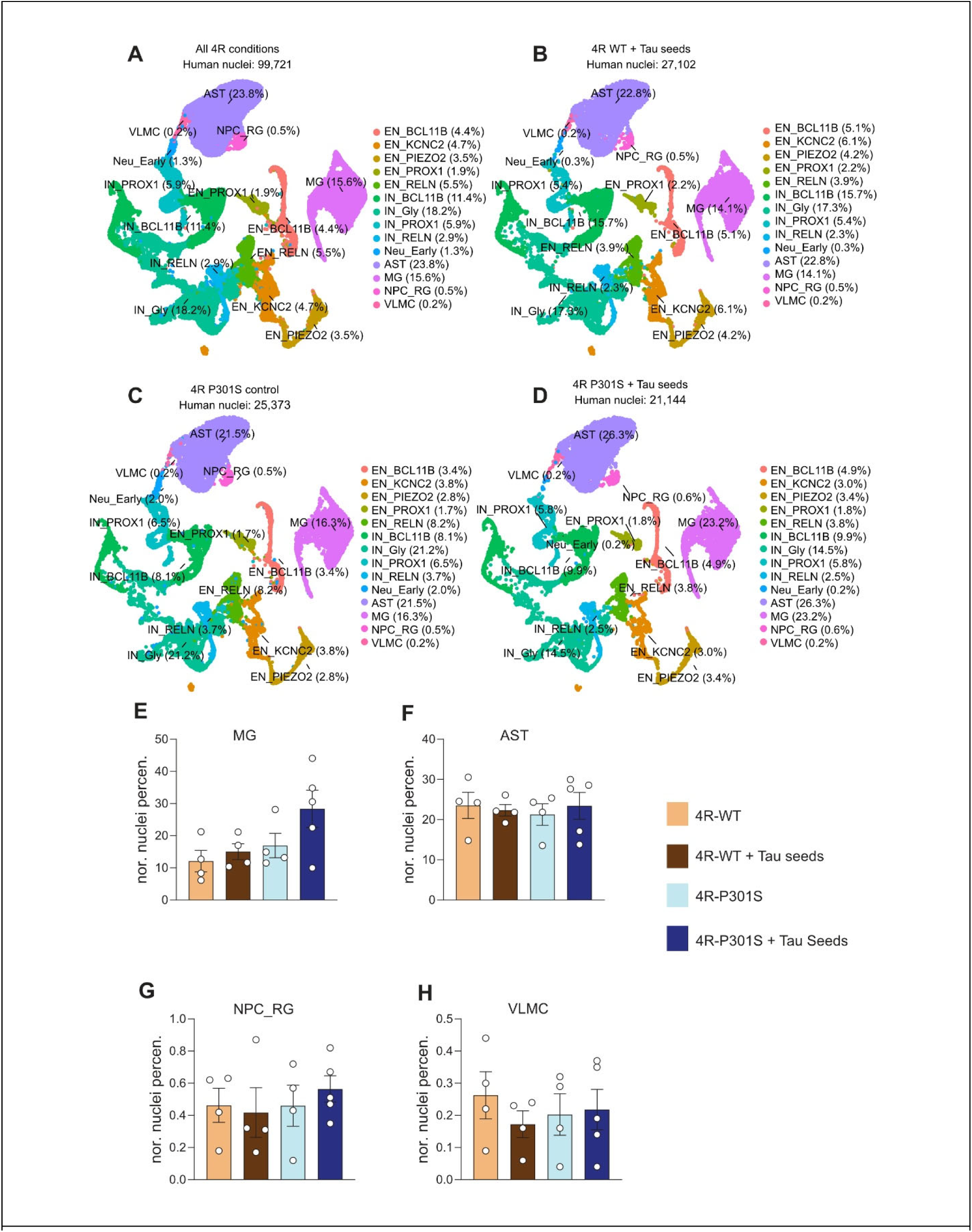
Map cellular populations in human grafts across 4R tau-seeding conditions using cell identities defined in 4R-WT grafts. **(A)** UMAP visualization of all human nuclei from the 4R conditions after transfer of 4R-WT cell type labels. Major human graft-derived populations and neuronal subtypes are annotated, with the percentage of total human nuclei indicated for each cluster. (**B-D**) Condition-specific UMAP visualizations showing transferred cell type labels in 4R-WT plus tau seeds (B), 4R-P301S control (C), and 4R-P301S plus tau seeds (D) grafts. The number of recovered human nuclei is indicated above each UMAP. (**E-H**) Quantification of the relative abundance of non-neuronal human graft populations across conditions, including human microglia, MG (E), astrocytes, AST (F), neural progenitor or radial glia-like cells, NPC_RG (G), and vascular leptomeningeal-like cells, VLMC (H). Data are presented as mean ± SEM. Each dot represents one mouse. Condition colors indicate 4R-WT, 4R-WT plus tau seeds, 4R-P301S control, and 4R-P301S plus tau seeds.

**Fig. S8.**
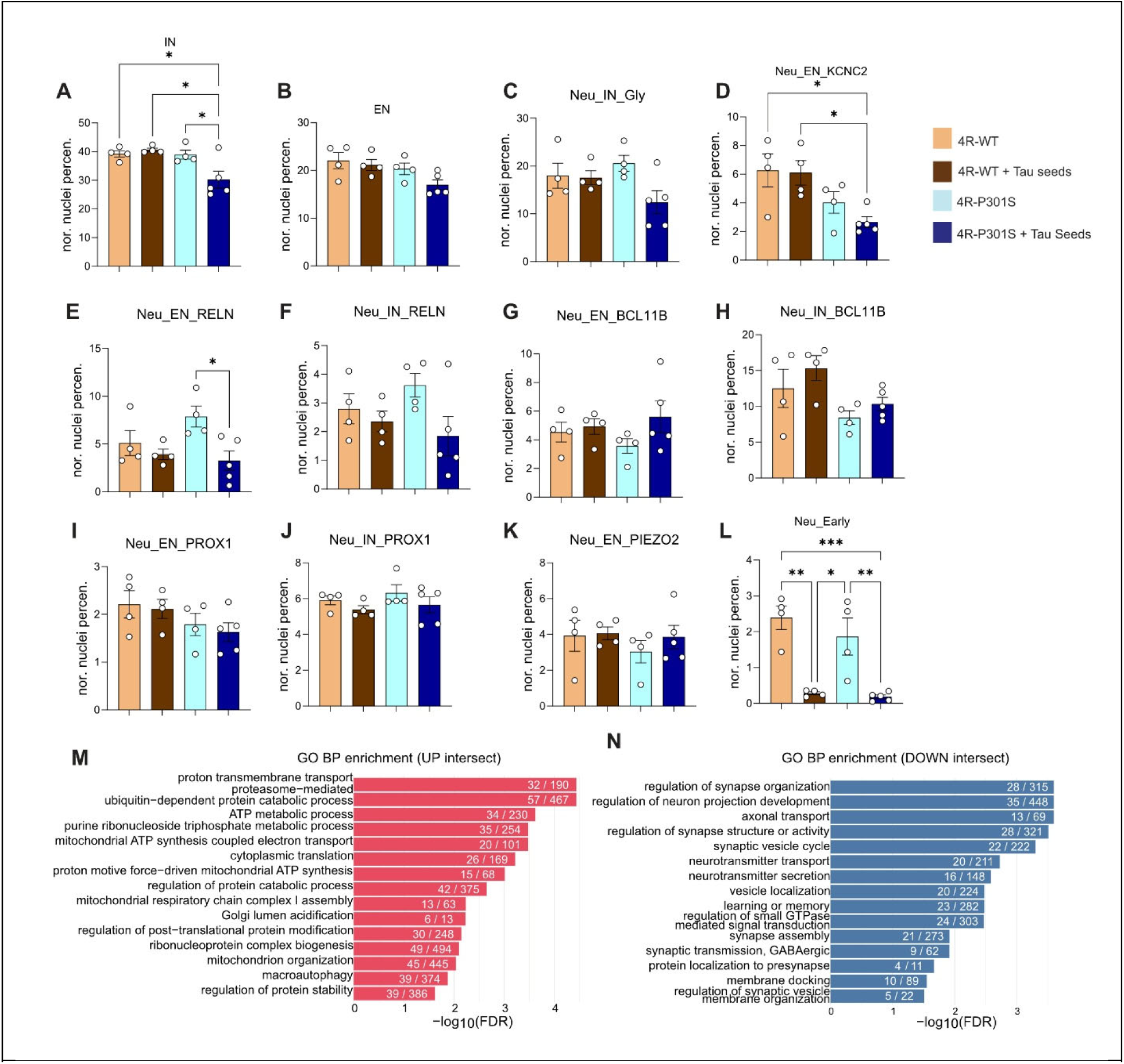
Relative abundance of human neuronal subtypes and shared pathway changes in vulnerable neuronal populations. (**A-L**) Quantification of human neuronal subtype abundance across 4R-WT, 4R-WT plus tau seeds, 4R-P301S control, and 4R-P301S plus tau seeds grafts. Bar plots show the proportional representation of major inhibitory neurons, IN (A), excitatory neurons, EN (B), glycinergic inhibitory neurons (C), KCNC2-high excitatory neurons (D), RELN-high excitatory neurons (E), RELN-high inhibitory neurons (F), BCL11B-high excitatory neurons (G), BCL11B-high inhibitory neurons (H), PROX1-high excitatory neurons (I), PROX1-high inhibitory neurons (J), PIEZO2-high excitatory neurons (K), and early neurons (L). (**M, N**) Gene Ontology biological process enrichment analysis of commonly dysregulated genes shared by vulnerable neuronal populations identified in Fig. 3, including SLC17A6-high, SLC6A5-high, KCNC2-high, and RELN-high neuronal populations. Enrichment results are shown for commonly upregulated genes (M) and commonly downregulated genes (N) in tau-seeded 4R-P301S grafts. Numbers within bars indicate the number of overlapping genes and the total number of genes in each GO term. Data are presented as mean ± SEM. Each dot represents one mouse. N = 4 to 5 mice per condition. Statistical significance was determined by one-way ANOVA followed by Tukey’s post hoc test. *P < 0.05, **P < 0.01, ***P < 0.001.

**Fig. S9.**
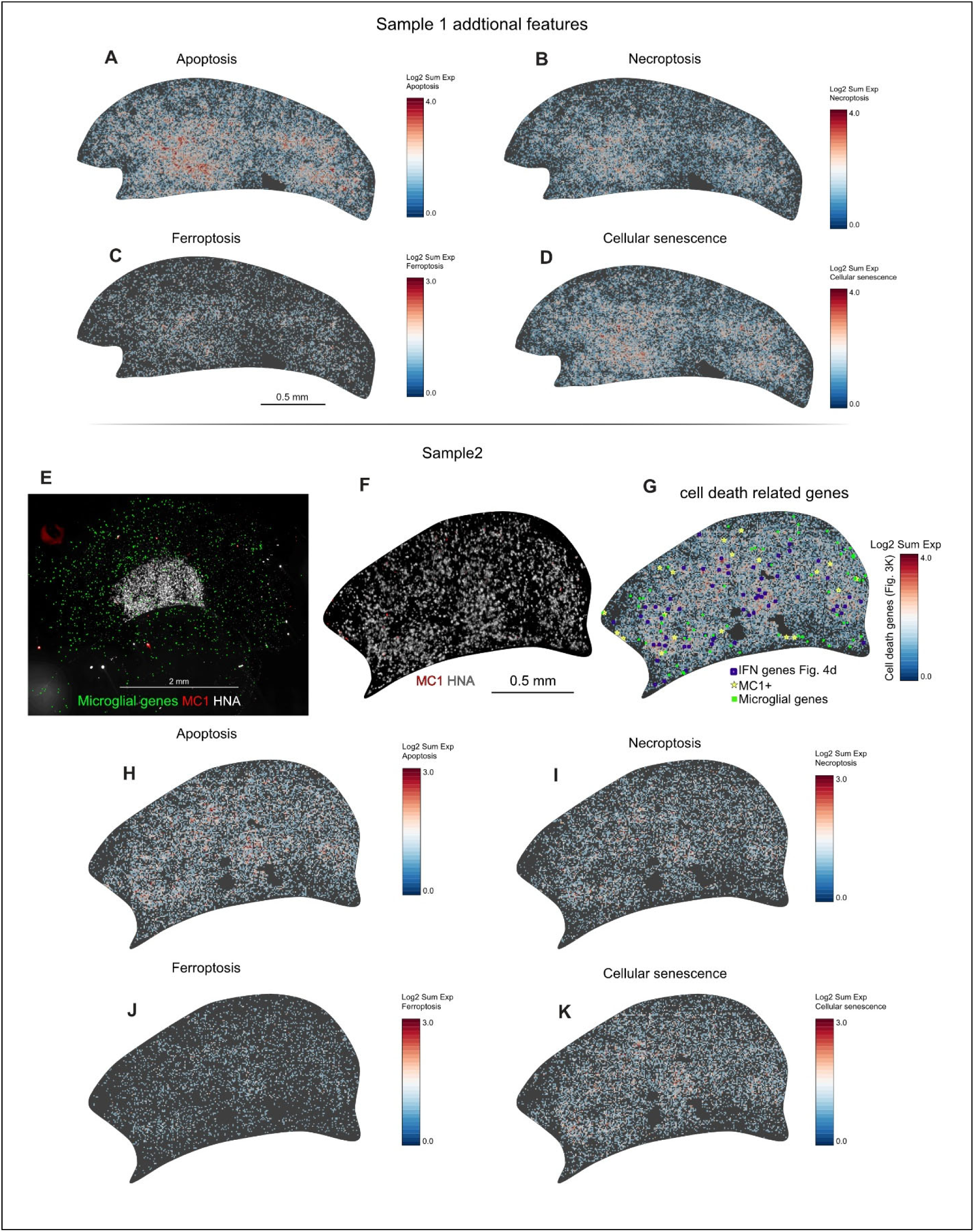
Visium HD spatial mapping of cell death related transcriptional programs and human microglial distribution. (**A-D**) Visium HD spatial transcriptomic maps from Sample 1 showing spatial enrichment of curated gene sets related to apoptosis (A), necroptosis (B), ferroptosis (C), and cellular senescence (D). Spatial expression is shown as log2 summed expression for each gene set. (**E-G**) Independent Visium HD dataset from Sample 2 showing the spatial relationship among human microglia, tau pathology, interferon associated regions, and cell death related transcriptional programs. E, Representative section showing human microglial gene expression, MC1 positive tau pathology, and human nuclei antigen within and around the graft. Microglial genes included CX3CR1, SALL1, CSF1R, P2RY12, P2RY13, SLCO2B1, TMEM119, and SPI1. F, Spatial localization of MC1 pathology and human nuclei antigen in Sample 2. G, Spatial map of the combined cell death related gene program identified in Fig. 3K, overlaid with IFN associated genes from Fig. 4D, MC1 positive regions, and human microglial gene expression. (**H-K**) Spatial enrichment of apoptosis (H), necroptosis (I), ferroptosis (J), and cellular senescence (K) gene sets in Sample 2. Together, these independent Visium HD datasets support spatial association among tau pathology, human microglial distribution, interferon associated signals, and cell death related transcriptional programs in HuMiNAX grafts. Gene sets used for spatial scoring were derived from the pathway analyses in Fig. 3K.

**Fig. S10.**
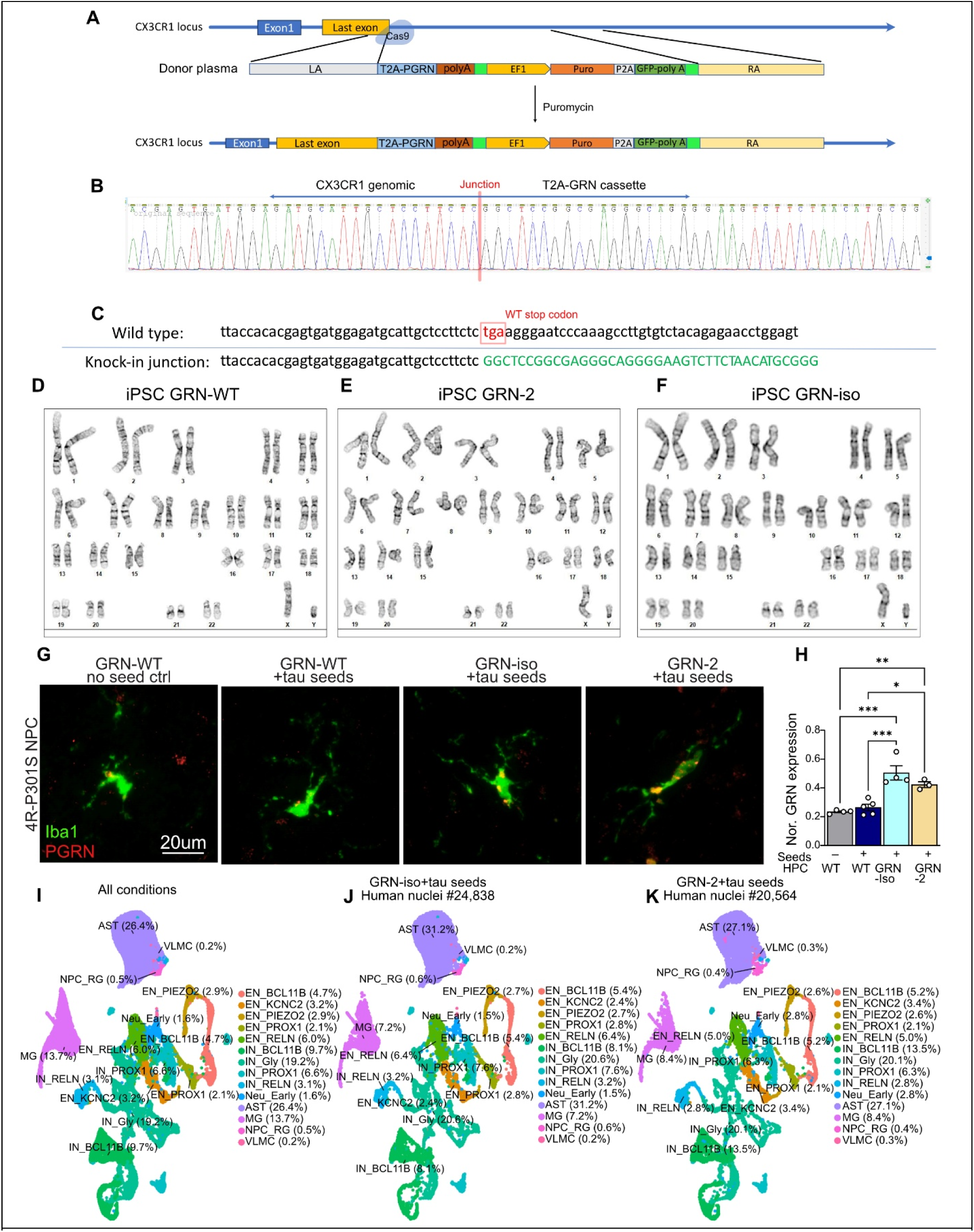
Generation and validation of CX3CR1-driven microglial GRN-overexpression iPSC lines. (**A**) CRISPR-Cas9-mediated homology-directed knock-in strategy at the human CX3CR1 locus. A guide RNA cut site was positioned near the CX3CR1 stop codon to enable in-frame insertion of a donor cassette containing T2A-PGRN, followed by an EF1α-puromycin-P2A-GFP selection module and flanked by CX3CR1 left and right homology arms. Correctly targeted clones were enriched by puromycin selection. Schematic is not to scale. (**B**) Sanger sequencing chromatogram of a junction PCR amplicon spanning the CX3CR1 genomic sequence and the inserted T2A-GRN cassette. The red tick marks the knock-in junction, confirming targeted integration at the intended locus. (**C**) Sequence alignment of wild-type and knock-in alleles showing replacement of the endogenous CX3CR1 stop codon with the expected T2A-GRN cassette sequence. (**D-F**) Representative G-band karyotypes showing normal chromosomal complements in the parental GRN-WT iPSC line (D) and two independent CX3CR1-T2A-GRN knock-in iPSC clones, GRN-2 (E) and GRN-iso (F). (**G**) Representative immunostaining for Iba1 and PGRN in HuMiNAX grafts containing GRN-WT microglia without tau seeds, GRN-WT microglia with tau seeds, GRN-iso microglia with tau seeds, or GRN-2 microglia with tau seeds. PGRN-overexpressing microglia showed increased PGRN signal in graft-resident human microglia compared with GRN-WT controls. (**H**) Sample-level quantification of normalized aggregate GRN expression in graft-derived microglia from the snRNA-seq dataset. GRN expression was increased in grafts containing GRN-iso or GRN-2 microglia compared with tau-seeded GRN-WT controls. No-seed WT-HPC, n = 4 mice; seeded WT-HPC, n = 5 mice; GRN-iso, n = 4 mice; GRN-2, n = 3 mice. One-way ANOVA followed by Tukey’s post hoc test. *P < 0.05, **P < 0.01, ***P < 0.001, ****P < 0.0001. (**I**) UMAP visualization of the re-integrated snRNA-seq dataset containing WT-HPC non-seeded and tau-seeded reference samples together with PGRN-OE tau-seeded samples. Major graft-derived human cell populations and neuronal subtypes are annotated, with percentages indicating the proportion of each annotated population among human nuclei. (**J-K**) UMAP visualization of individual PGRN-OE conditions from the re-integrated dataset, showing annotated human nuclei from GRN-iso plus tau seeds (J) and GRN-2 plus tau seeds (K).

**Fig. S11.**
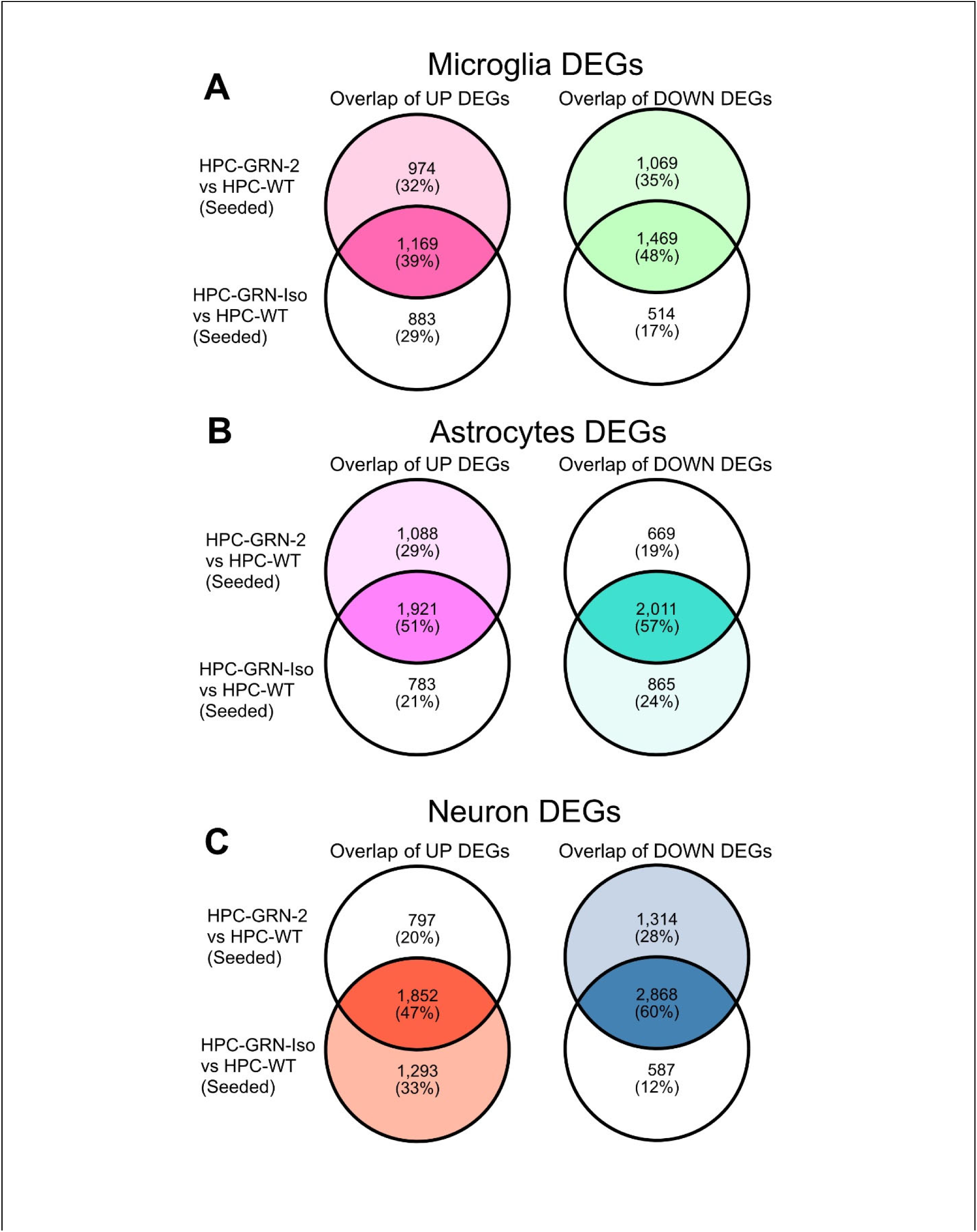
Consensus transcriptional effects of two independent PGRN-overexpressing microglial lines across human graft cell types. (**A-C**) Venn diagrams showing the overlap of upregulated and downregulated DEGs induced by two independent PGRN-overexpressing microglial lines, GRN-2 and GRN-iso, relative to tau-seeded GRN-WT controls. Analyses are shown separately for graft-derived human microglia (A), astrocytes (B), and neurons (C). Percentages indicate the fraction of DEGs represented by each overlap category.

**Fig. S12.**
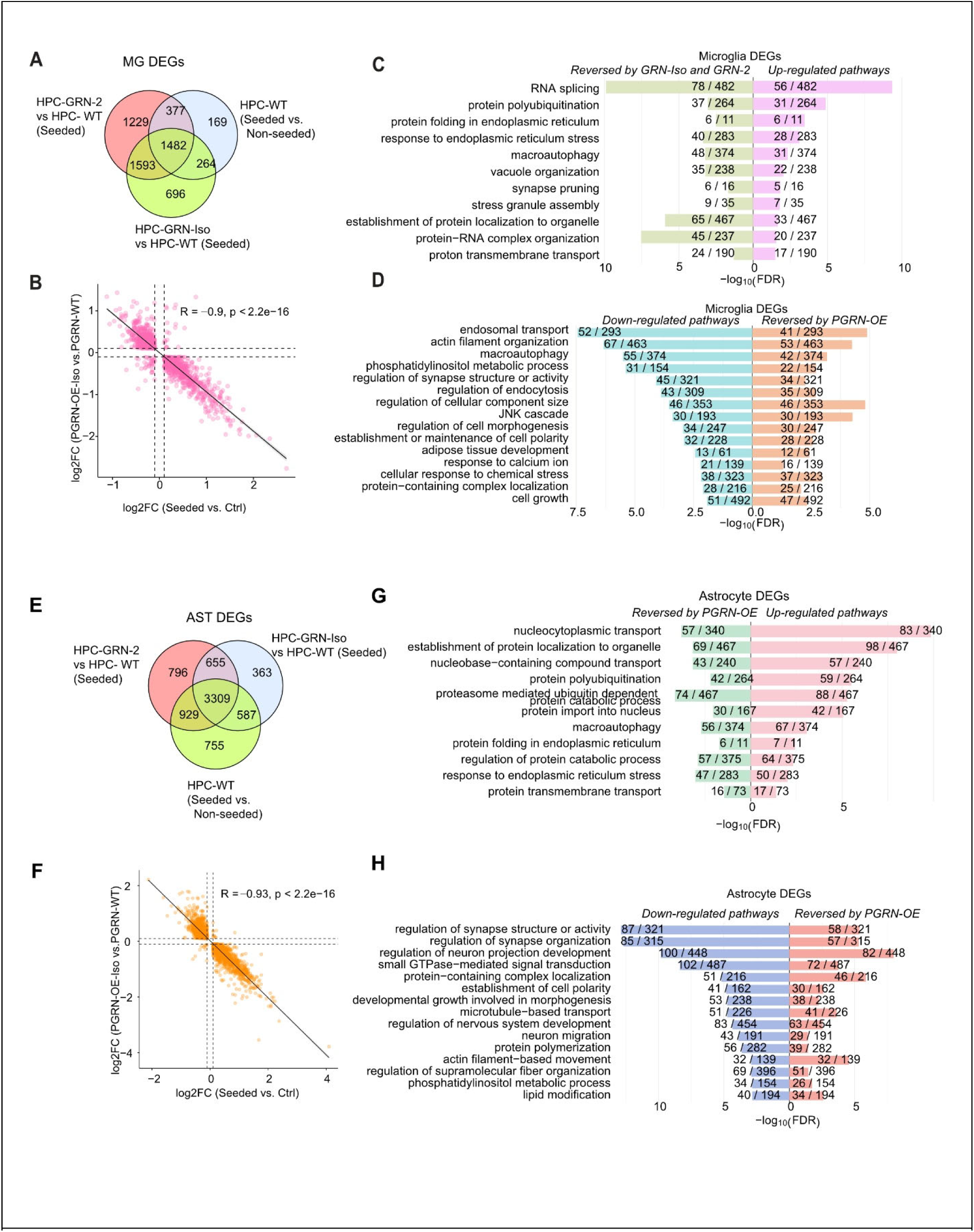
Microglial progranulin reverses tau-associated glial transcriptional programs in HuMiNAX. (**A-D**) Microglial DEG analysis. A, Venn diagram showing overlap among tau seed-associated microglial DEGs in WT-HPC grafts and DEGs induced by GRN-iso or GRN-2 microglia under tau-seeded conditions. B, Correlation of microglial log₂ fold changes showing inverse regulation between tau seed-induced changes and GRN-iso-associated changes. Pathway analyses showing tau-upregulated microglial programs reversed by GRN-iso and GRN-2 (C) and tau-downregulated programs restored by PGRN-OE microglia (D). (**E-H**) Astrocyte DEG analysis. E, Venn diagram showing overlap among tau seed-associated astrocyte DEGs in WT-HPC grafts and DEGs induced by GRN-iso or GRN-2 microglia under tau-seeded conditions. F, Correlation of astrocyte logfold changes showing broad reversal of tau-associated astrocyte changes by GRN-iso microglia. G,H, Pathway analyses showing tau-upregulated astrocyte programs reversed by PGRN-OE microglia (G) and tau-downregulated programs restored by PGRN-OE microglia (H). FDR, false discovery rate.

**Fig. S13.**
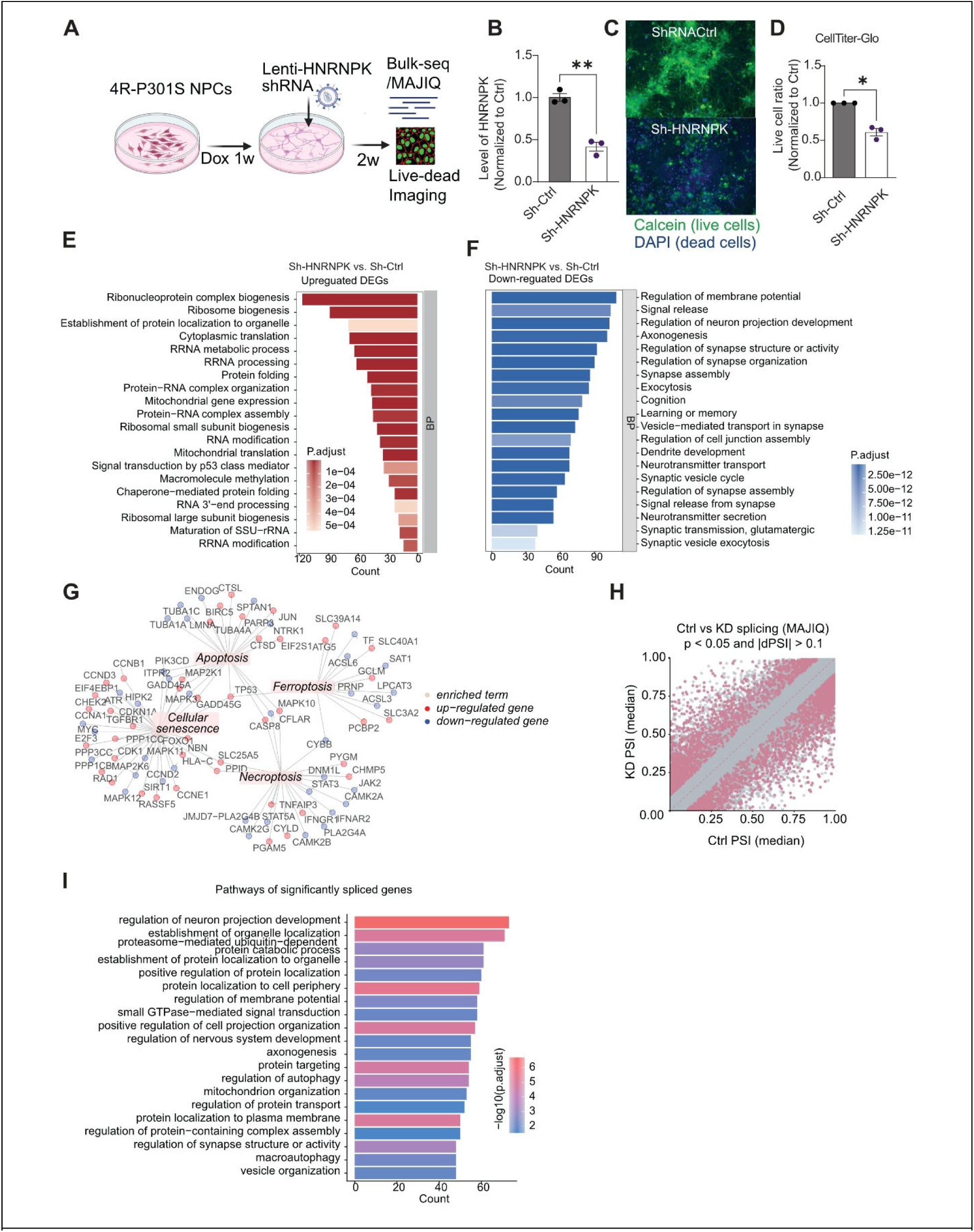
HNRNPK knockdown impairs survival and disrupts transcriptomic and splicing programs in human 4R P301S neurons. (**A**) Experimental schematic of lentiviral shRNA mediated HNRNPK knockdown in doxycycline induced NGN2 4R P301S human neurons. After neuronal induction, cells were transduced with control or HNRNPK targeting shRNA and analyzed by bulk RNA sequencing, MAJIQ splicing analysis, and live dead viability assays. (**B**) RT qPCR validation of HNRNPK knockdown in sh-HNRNPK neurons compared with sh control neurons. (**C-D**) Representative live dead images and CellTiter Glo quantification showing reduced neuronal viability following HNRNPK knockdown. Calcein marks live cells and DAPI marks dead cells. (**E-F**) Gene Ontology biological process enrichment analysis of differentially expressed genes following HNRNPK knockdown. Upregulated genes were enriched for ribonucleoprotein complex biogenesis, ribosome biogenesis, cytoplasmic translation, rRNA processing, protein folding, mitochondrial gene expression, and RNA modification pathways. Downregulated genes were enriched for regulation of membrane potential, synaptic signaling, neuron projection development, axonogenesis, synapse organization, neurotransmitter transport, and vesicle mediated synaptic processes. (**G**) Network visualization of cell death associated enriched terms and contributing differentially expressed genes following HNRNPK knockdown, highlighting apoptosis, ferroptosis, cellular senescence, and necroptosis related gene programs. (**H**) MAJIQ analysis of alternative splicing changes between sh control and sh HNRNPK neurons. Scatter plot shows median percent spliced in values for control and HNRNPK knockdown conditions. Significantly altered local splicing variations were defined by t-test P < 0.05 and |dPSI| > 0.1. (**I**) Gene Ontology biological process enrichment analysis of genes with significantly altered splicing events following HNRNPK knockdown. Data are shown as mean ± SEM. Each dot represents one independent differentiation and transduction. Statistical significance in B and D was determined by two tailed Welch’s t test. *P < 0.05, **P < 0.01.

**Fig. S14.**
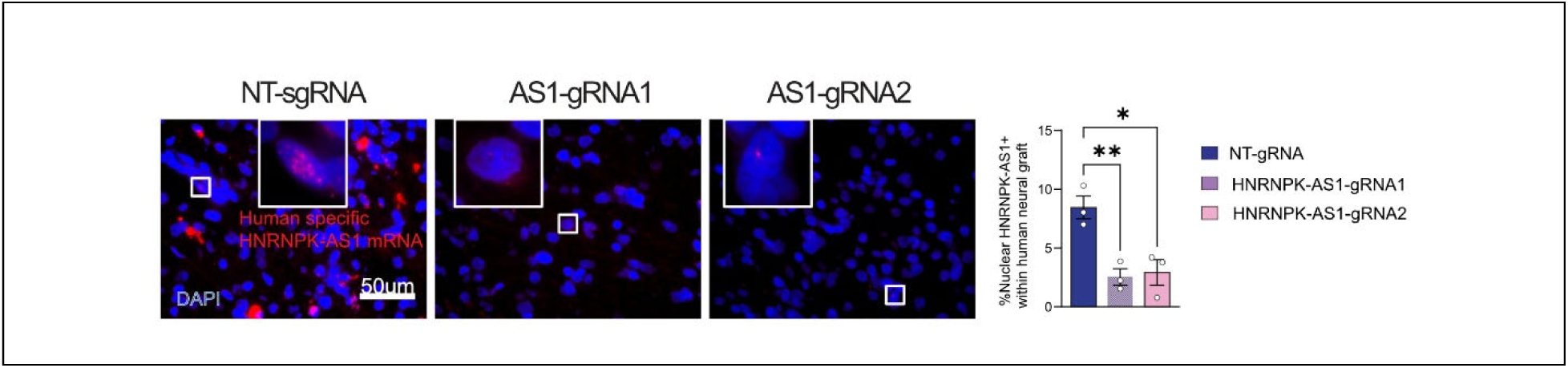
Validation of HNRNPK-AS1 knockdown in vivo. RNAscope in situ hybridization for human specific HNRNPK-AS1 in human neural grafts expressing non targeting sgRNA or two independent HNRNPK-AS1 targeting sgRNAs. Representative images and quantification show reduced nuclear HNRNPK-AS1 puncta within human nuclei following targeting with either HNRNPK-AS1 sgRNA. Insets show higher magnification views of boxed cells. Data are shown as mean ± SEM. Each dot represents one quantified graft. Statistical significance was determined by ordinary one way ANOVA followed by Tukey’s post hoc multiple comparisons test. *P < 0.05, **P < 0.01.

